# Class II Kinesin-12 facilitates cell plate formation by transporting cell plate materials in the phragmoplast

**DOI:** 10.1101/2024.08.20.608524

**Authors:** Moé Yamada, Hironori J. Matsuyama

**Affiliations:** Division of Biological Science, Graduate School of Science, Nagoya University, Nagoya, Japan; Neuroscience Institute, Graduate School of Science, Nagoya University, Nagoya, Japan

## Abstract

Cell plate formation in plants is a complex process orchestrated by the targeted delivery of Golgi-derived and endosomal vesicles containing cell plate components to the phragmoplast midzone. It has long been hypothesised that vesicles are directionally transported along phragmoplast microtubules by motor proteins. However, the mechanisms governing the accumulation and immobilisation of vesicles at the phragmoplast midzone remain elusive, and the motor protein responsible has yet to be identified. Here, we show that the plant-specific class II Kinesin-12 (Kinesin12-II) functions as a motor protein that drives vesicle transport towards the phragmoplast midzone in the moss *Physcomitrium patens*. In *kinesin12-II* mutants, the directional movement of cell plate materials towards the midzone and their retention were abolished, resulting in delayed cell plate formation and phragmoplast disassembly. A macroscopic phenotype arising from *Kinesin12-II* disruption was the impediment to gametophore development. We showed that this defect was attributable to the production of aneuploid cells in the early gametophore, where chromosome missegregation occurred because of the incomplete disassembly of phragmoplast microtubules in the preceding cell cycle. These findings suggest that plant Kinesin-12 has evolved to acquire a unique and critical function that facilitates cell plate formation in the presence of phragmoplasts.

## Introduction

Cytokinesis is the final step in cell division in which cellular contents are partitioned into two daughter cells. In plants, the mother cell is partitioned by the formation of the cell plate between two daughter nuclei, which centrifugally expands towards the cell cortex from the centre of the cell ^1^. As the cross-wall and positional relationships between two daughter cells are permanently maintained in plants, the insertion of a new cell wall at an accurate position is particularly important for development and morphogenesis. Cell plate formation occurs in the midzone of the phragmoplast, which is a cytoskeletal scaffold consisting of two opposing sets of microtubule arrays, actin filaments and membranes ^2–4^. The mechanism by which phragmoplast microtubules are assembled, maintained and disassembled during cytokinesis has been extensively studied, and key molecules have been identified, such as MAP65 which cross-links antiparallel microtubules at the midzone ^5–14^. The mechanism of the final step of cytokinesis, namely the remodelling of cell plate materials ^1^, is also fairly well understood. The vesicles which accumulate at the midzone are fused and sequentially remodelled into a tubule–vesicular network, tubular network and planar fenestrated sheet; this is orchestrated by clathrin-mediated endocytosis, and the sheet then forms a cross-wall ^4,15,17^. Genetic approaches using *Arabidopsis* have identified factors responsible for vesicle fusion and cell plate maturation, such as KNOLLE, RABA2, TRAPII and the Exocyst complex ^18–21^. However, little is known about the recruitment and immobilisation of cell plate materials at the phragmoplast midzone.

Cell plate formation begins with the accumulation of Golgi-derived and endosomal vesicles at the phragmoplast midzone. They constitute the cell plate assembly matrix (CPAM) together with polysaccharide synthase and vesicle-associated molecular machinery^1,18,22^. The mechanisms of precise targeting and entrapment of vesicles in the phragmoplast midzone are not yet understood at the molecular level. A prevailing model hypothesises that vesicles are directionally transported along phragmoplast microtubules by motor proteins. This model was proposed based on two electron microscopy observations: 1) many vesicles are positioned close to microtubules in the phragmoplast ^4,23^; 2) a kinesin-like ultrastructure that is associated with the vesicle as well as microtubules was detected ^24^. However, whether the vesicles are actually moved along the phragmoplast microtubules towards the midzone is not known. The identity of the detected kinesin-like motor and whether the motor transports cell plate materials have not been revealed. Furthermore, it is intuitively unclear why plant cells deploy an energy-consuming transport mechanism rather than diffusion for cytokinetic vesicle accumulation—the phragmoplast (∼10 µm) is not a large or enclosed structure compared to extremely long axons or diffusion-limited cilia wherein directional transport would be much more favourable than diffusion for protein/vesicle localisation at the tip ^25^. Thus, the mechanism by which vesicles accumulate and are immobilised at the phragmoplast midzone remains unknown.

Kinesin-12 is a conserved kinesin subfamily ^26^. Mammalian KIF15/Kinesin-12 drives the outwards sliding of antiparallel microtubules in the spindle, contributing to the maintenance of spindle bipolarity^27,28^. Land plant Kinesin-12 subfamily comprises two classes (I and II), with 6 and 20 members in *Arabidopsis* and the moss *Physcomitrium patens*, respectively ^29,30^. Despite its wide conservation in plants, most of our knowledge of plant Kinesin-12 is based on studies on *Arabidopsis*. *Arabidopsis* class I Kinesin-12 (Kinesin12-I, POK1/Kin12C and POK2/Kin12D) mediates accurate cell plate positioning by guiding the expanding phragmoplast to predetermined cortical division sites ^31^. Class II Kinesin-12 (Kinesin12-II) is comprised of three members in *Arabidopsis*: PAKRP1/Kin12A, PAKRP1L/Kin12B and Kin12F. Kin12A and Kin12B are localised at the phragmoplast midzone^32^. The *kin12a kin12b* double mutant exhibits phragmoplast disorganisation and failure of cell plate formation during cytokinesis^33^. It has been proposed that Kin12A and Kin12B play a critical role in allowing phragmoplast microtubules to be organised into two mirrored sets, with a gap between them.

The moss *P. patens* is an excellent model system for studying cytokinesis because it is amenable to microscopic observation without the need for dissection or sectioning, owing to its small stature and anatomical simplicity ^34^. Four *Kinesin12-II* genes are encoded in *P. patens*. However, their functions have not been investigated. In this study, we aimed to test the long-standing hypothesis that vesicular transport by motor proteins plays a critical role in cell plate formation, by using the protonemal tissue of *P. patens*. We provide evidence that cytokinetic vesicles are transported directionally along phragmoplast microtubules. We then identified Kinesin12-II as the driver of cell plate material transport towards the phragmoplast midzone. Furthermore, we elucidated the critical role of transport-mediated cell plate formation in moss development through live observation of *kinesin12-II* mutants.

## Results

### Endogenous SCAMP4-tagging visualises the directional movement of cell plate materials in the phragmoplast

To investigate how cell plate materials accumulate at the phragmoplast midzone during cytokinesis, we visualised the movement of vesicles in the phragmoplast by endogenously tagging secretory carrier membrane protein 4 (SCAMP4) with the green fluorescent protein mNeonGreen (mNG) in the moss *P. patens* ^10,35^. During cytokinesis in protonemal cells, punctate SCAMP4–mNG signals were detected in both the cytoplasm and phragmoplasts. In addition, a linear signal corresponding to the cell plate appeared at the midzone after anaphase onset (Fig. 1a). Time-lapse imaging with high-resolution microscopy indicated that SCAMP4–mNG puncta exhibited both directional and diffusive motion in the cell. Interestingly, some punctate signals in the phragmoplast moved directionally towards the cell plate, suggesting that the accumulation of cell plate materials at the midzone involves a transport-mediated mechanism (Fig. 1b and Supplementary Video 1). We measured the speed of the directional motion and found that the velocity of SCAMP4–mNG signals in the phragmoplast and cytoplasm showed different distribution patterns; motility > 1.0 µm/s was observed only in the cytoplasm (Fig. 1c, d). This observation suggests that multiple mechanisms drive the subcellular movements of SCAMP4-marked vesicles and that cell plate-directed movement is regulated in a manner distinct from that in the cytoplasm.

**Figure 1.**
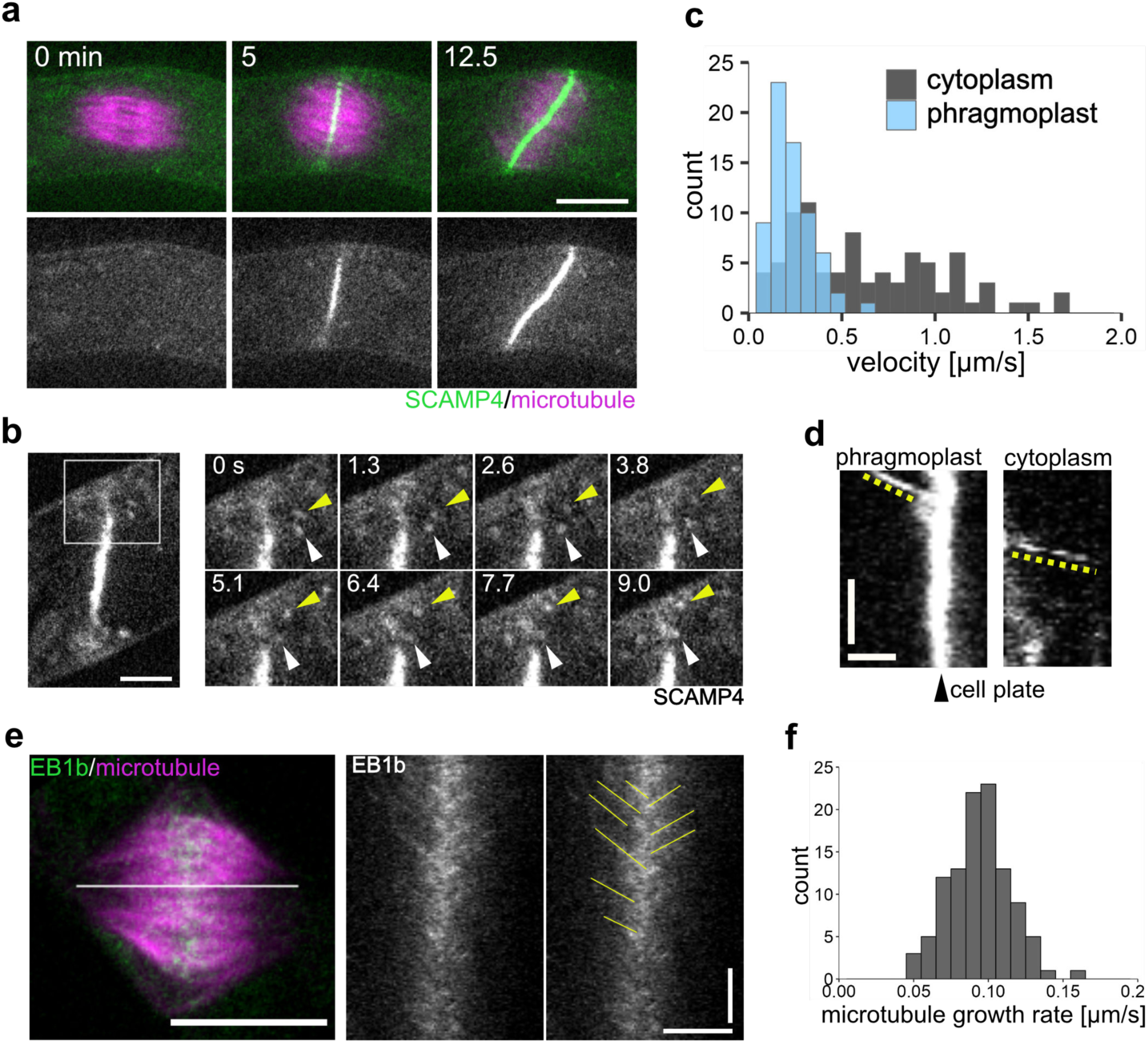
SCAMP4–mNG marker visualises the directional motility of cell plate materials in the phragmoplast. (a) Snapshots of a dividing protonemal tip cell expressing mCherry–α-tubulin (magenta) and SCAMP4– mNG (green). A single plane of confocal images is presented. Time 0 corresponds to anaphase onset. Chloroplast autofluorescence is also faintly visualised in the green channel. Scale bar, 10 µm. (b) Snapshots of a protonemal tip cell expressing SCMAP4–mNG during cytokinesis. A single plane of confocal images is presented. The right panels are magnified images of the area indicated with the square bracket in the left panel. Arrowheads indicate punctate signals that exhibit directional movement towards the leading edge of the phragmoplast. Scale bar, 5 µm. (c) Quantitative analysis of SCMAP4 signal velocity in the phragmoplast (blue) and the cytoplasm (dark grey). Phragmoplast, 0.23 ± 0.11 µm/s (mean ± SD), n = 68; cytoplasm, 0.63 ± 0.40 µm/s, n = 83. N > 10 cells were analysed for each quantification. (d) Kymographs showing directional movements of SCAMP4 signals in the phragmoplast and the cytoplasm. Directional motility is represented by oblique lines in the kymographs. The black arrowhead indicates the position of the cell plate, and yellow dashed lines indicate the path of the directional movements. Horizontal bar, 2 µm; vertical bar, 20 s. (e) Snapshots of the phragmoplast in a dividing protonemal tip cell expressing mCherry–α-tubulin (magenta) and EB1b–mNG (green). A single plane of confocal images is presented. The right panels are kymographs of EB1b–mNG, generated from the region indicated by a white line in the left panel. In the kymographs, yellow lines highlight directional movement trajectories of the EB1b–mNG signals. Scale Bar (left), 10 µm; Horizontal bar (right), 5 µm; vertical bar (right), 1 min. (f) Histogram depicting microtubule growth rate velocity measured by the dynamics of EB1b–mNG. 0.09 ± 0.02 µm/s (mean ± SD), n = 107.

Both actin- and microtubule-based motility play a role in intracellular transport in plants, whereas actin-based motility is dispensable for cell plate formation in several cell types^36–40^. We confirmed in our system that actin inhibition by latrunculin A treatment did not perturb cell plate formation; the linear signals of SCAMP4–mNG appeared normal, and phragmoplast microtubules disassembled without a significant delay upon latrunculin A treatment (Supplementary Fig. 1a–c). We concluded that actin was not involved in the transport of SCAMP4-marked vesicles.

These results suggest that microtubule-based motility drives the transport of SCAMP4-marked vesicles in the phragmoplast. Microtubule polymerisation is a possible driver of intracellular motility as exemplified by ER tubule protrusion in animal cells, in which the ER protein STIM1 binds to the tip of polymerising microtubules via the microtubule plus-end tracking protein EB1^41,42^. To test the validity of this model in moss phragmoplasts, we measured the polymerisation rate of phragmoplast microtubules by observing endogenously tagged EB1b ^43^. The majority of EB1b–mNG signals moved towards the midzone at an average of 94 ± 20 nm/s (mean ± SD, n = 107) (Fig. 1e, f), which was two-fold slower than the SCAMP4 vesicles. Thus, microtubule polymerisation is unlikely to be the main driver of vesicle motility.

### Kinesin12-II affects protonemal growth and gametophore development

We considered the possible contribution of kinesin superfamily proteins to vesicular transport, as kinesin is a microtubule-based motor protein associated with various cellular processes, including intracellular transport ^26,44^. To identify the specific kinesin gene responsible for vesicle transport towards the cell plate in the phragmoplast, we used a reverse genetic approach. We reasoned that a motor protein transporting vesicles towards the cell plate in the phragmoplast would meet the following criteria: 1) it would have plus-end-directed motility, as the microtubule plus-ends face the cell plate in the phragmoplast; 2) it would be localised at the phragmoplast midzone or along the phragmoplast microtubules; 3) its cellular function has not been revealed; alternatively, the loss-of-function phenotype has been shown to be cytokinesis-related but has not been analysed in the context of vesicle transport. Among the 78 kinesin proteins present in *P. patens*, we suspected that the Kinesin12-II subfamily could be the strongest candidate transporter that would fulfil all three criteria.

Four *Kinesin12-II* genes are encoded in the genome of *P. patens* and are referred to herein as *Kinesin12-IIa, -IIb, -IIc* and *-IId* ^11,30^. A previous study showed phragmoplast midzone localisation in three of these following citrine tagging ^11^. In the present study, we inserted the *mNG* gene into the endogenous locus via homologous recombination and observed that all four Kinesin12-IIs showed strong enrichment at the phragmoplast midzone and faint localisation in the entire phragmoplast (Supplementary Fig. 2). Next, we generated *kinesin12-II* quadruple mutants by introducing mutations into all four *Kinesin12-II* genes using CRISPR/Cas9-mediated genome editing ^45,46^. To eliminate motor activity, we targeted the SSRSH and LAGSE motifs, which consist of Switch 1 and Switch 2 loops, respectively, in the kinesin motor domain, and are indispensable for ATP hydrolysis ^28^. In the background of mCherry–α-tubulin or GFP–α-tubulin/histoneH2B–mCherry expression lines, we obtained multiple independent lines that possessed various mutations in different gene sets (Fig. 2a, b). Among them were two quadruple mutant lines, which carried frameshift mutations in the *Kinesin12-IIa, -IIb* and *-IId* genes and one amino acid deletion in the *Kinesin12-IIc* gene (hereafter referred to as *kin12-IIabcd-1* and *kin12-IIabcd-2* alleles) (Supplementary Fig. 3). We could not obtain a line in which a frameshift was introduced in all four genes after several attempts, suggesting that the line completely null for Kinesin12-II was lethal. Disruption of *Kinesin12-II* genes caused severe defects in protonemal growth and gametophore (leafy shoot) development; plant sizes were substantially smaller than the control and no mature gametophores were produced on the culture plate (Fig. 2a, b). Growth defects in protonemal cells and gametophore developmental disorders were restored by expressing Kinesin12-IIc– mNG under an endogenous promoter, suggesting the functionality of the fluorescently tagged Kinesin12-IIc protein and functional redundancy among Kinesin12-IIs (Fig. 2a, b). We concluded that Kinesin12-II plays important roles in plant growth and gametophore development in *P. patens*.

**Figure 2.**
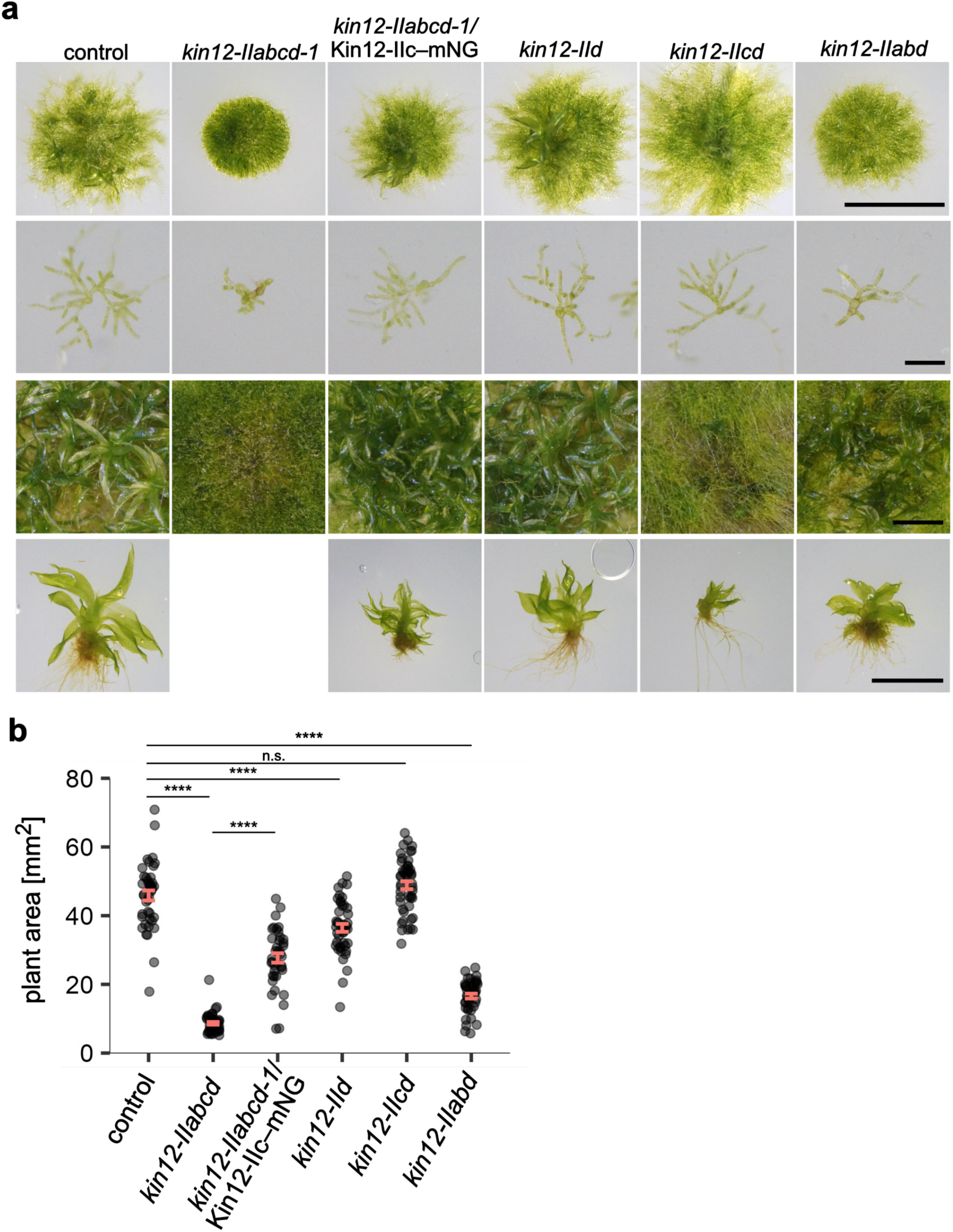
*Kinesin12-II* disruption causes growth retardation and gametophore developmental disorder. (a) Snapshots of three-week-old moss (top panels), one-week-old protonematal cells (second row panels), five-week-old moss (third row panels) and the gametophores (bottom panels). These moss samples were cultured from single protoplasts. Scale bar, 5 mm (top), 200 µm (second) and 2 mm (third and bottom). (b) Quantification of the moss size cultured for 3 weeks from protoplasts. Control, 45.9 ± 1.5 mm^2^ (mean ± SEM), n = 43; *kin12-IIabcd-1*, 8.7 ± 0.4 mm^2^, n = 49; *kin12-IIabcd-1*/Kin12-IIc-mNG, 27.7 ± 1.4 mm^2^, n = 39; *kin12-IIc*, 36.4± 1.2 mm^2^, n = 45; *kin12-IIcd*, 48.9 ± 1.1 mm^2^, n = 48; *kin12-IIabd*, 16.6 ± 0.7 mm^2^, n = 43. *P*-value was calculated using ART ANOVA and Tukey’s test; *P* < 0.0001 (control– *kin12-IIabcd-1*), *P* < 0.0001 (*kin12-IIabcd-1*– *kin12-IIabcd-1*/Kin12-IIc-mNG), *P* < 0.0001 (control– *kin12-IId*), *P* > 0.3 (control– *kin12-IIcd*), *P* < 0.0001 (control– *kin12-IIabd*).

### Kinesin12-II is required for rapid phragmoplast disassembly

We examined whether mitosis and cytokinesis were defective in the *kinesin12-II* mutants using high-resolution time-lapse imaging of GFP–tubulin and histoneH2B–mCherry. In the *kin12-IIabcd-2* mutant, the mitotic spindle and phragmoplast were assembled in a manner indistinguishable from that of the control; there was no significant difference in mitotic duration between the control and mutant lines (Fig. 3a, b and Supplementary Video 2). However, phragmoplasts did not expand rapidly towards the cell edges in the mutant—it took longer for the phragmoplast microtubules to disassemble (Fig. 3c). During prolonged phragmoplast expansion in the mutant, the phragmoplast occasionally rotated drastically and was displaced within the cell (Fig. 3a, d, e). Interestingly, the phragmoplast was disorganised during expansion in the mutant (Fig. 3a, 15 min). We quantified the GFP–tubulin intensity along the phragmoplast from one phragmoplast pole to the other. The intensity plot showed a single GFP–tubulin intensity peak at the midzone immediately after anaphase onset in both the control and *kin12-IIabcd-2* mutant, suggesting that antiparallel microtubule overlap formation occurred normally at the midzone in the mutant (Fig. 3f). However, although the GFP–tubulin intensity at the midzone was reduced 15 min after anaphase onset in the control, this was not the case in the *kin12-IIabcd-2* mutant (Fig. 3g). The lack of a microtubule gap in the mutant is reminiscent of the *kin12a kin12b* double mutant phenotype in *Arabidopsis* ^33^. To validate the formation of antiparallel microtubule overlaps in the phragmoplast, we used the MAP65 marker ^13,47^. Consistent with our interpretation, MAP65c–citrine signals were detected at the phragmoplast midzone in the *kin12-IIabcd-1* mutant, similar to in the control (Fig. 3h, i and Supplementary Video 3). These results suggest that moss Kinesin12-II is not required for the formation of antiparallel microtubule overlaps. Next, we investigated whether proper microtubule polarity was maintained in the phragmoplast by observing the dynamics of EB1b–mNG. Most EB1b–mNG signals moved towards the midzone in the phragmoplast of the mutant, indicating that the overall microtubule polarity was not perturbed (Supplementary Fig. 4a and Supplementary Video 4). Microtubule growth rate was also not significantly altered (Supplementary Fig. 4b). Taken together, we concluded that phragmoplast expansion and disassembly, but not phragmoplast formation, are defective in the *kinesin12-II* mutants.

**Figure 3.**
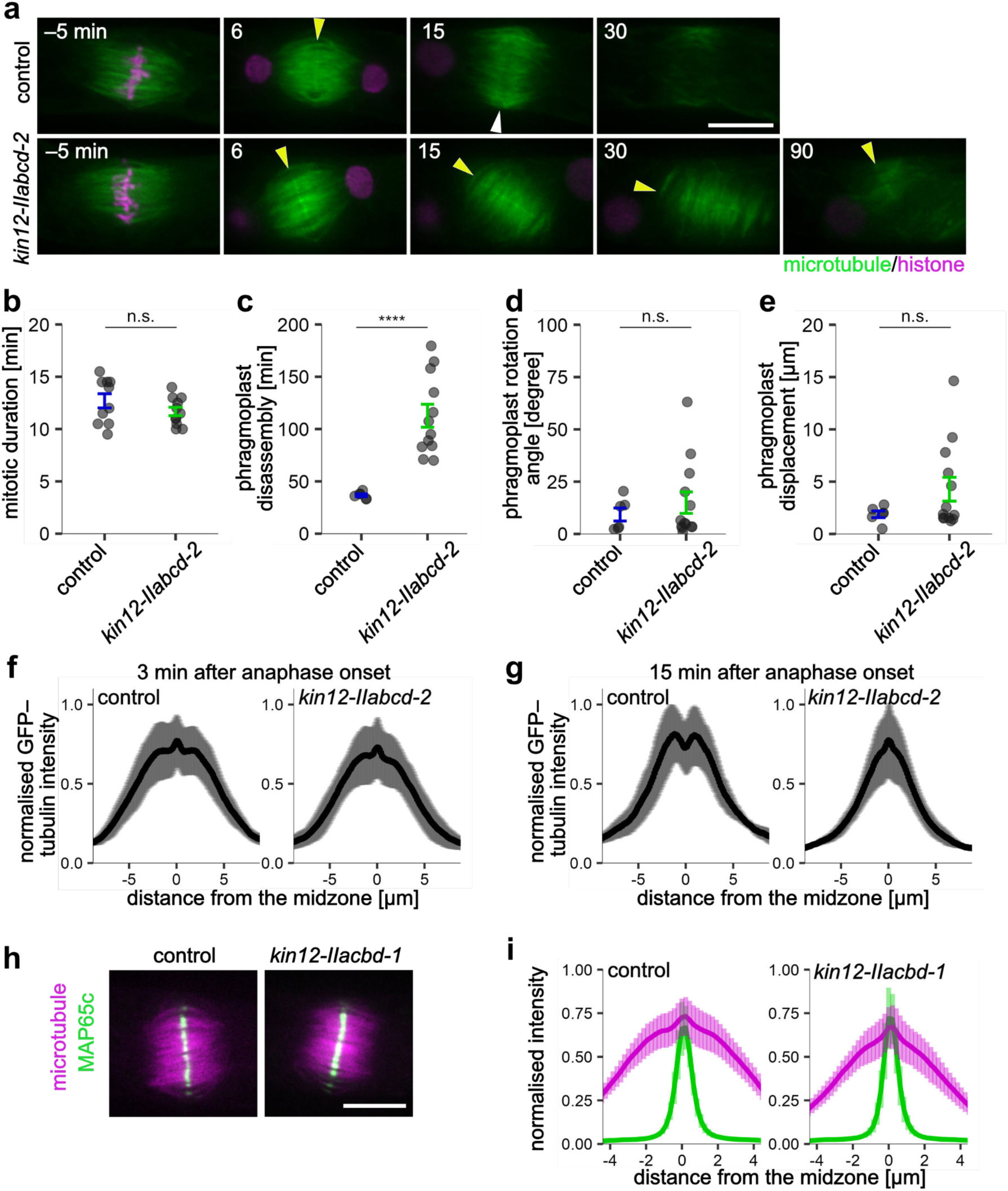
Phragmoplast formation is not affected by *Kinesin12-II* disruption. (a) Snapshots of a dividing protonemal tip cell expressing GFP–α-tubulin (green) and histoneH2B– mCherry (magenta). Maximum intensity projections derived from three confocal images acquired every 2 µm are presented. Yellow arrowheads indicate the GFP–α-tubulin peak at the phragmoplast midzone, and the white arrowhead indicates a gap formed between two opposing set of phragmoplast microtubules. Time 0 corresponds to anaphase onset. Scale bar, 10 µm. (b–e) Comparison of mitotic duration, phragmoplast disassembly duration, phragmoplast rotation angle and phragmoplast displacement. (b) Control, 12.7 ± 0.7 min (mean ± SEM), n = 10; *kin12-IIabcd-2*, 11.7 ± 0.4 min, n = 11. *P*-value was calculated using the Student’s *t*-test, *P* = 0.21; (c) control, 36.7 ± 1.3 min (mean ± SEM), n = 6; *kin12-IIabcd-2*, 112.7 ± 11.0 min, n = 12. *P*-value was calculated using the Welch’s *t*-test, *P* < 0.0001; (d) control, 9.3 ± 3.1 degrees (mean ± SEM), n = 6; *kin12-IIabcd-2*, 15.0 ± 5.1 degrees, n = 13. *P*-value was calculated using the Mann–Whitney *U* test, *P* > 0.5; (e) control, 1.9 ± 0.3 µm (mean ± SEM), n = 6; *kin12-IIabcd-2*, 4.3 ± 1.1 µm, n = 13. *P*-value was calculated using the Brunner–Munzel test, *P* > 0.6. (f–g) Normalised intensity profile of GFP–α-tubulin in the phragmoplast of the control line (left) and *kin12-IIabcd-2* mutant (right). The intensity was measured perpendicular to the division plane. The midzone position was plotted at 0 µm in the graphs. The mean intensity is depicted with a black line, and the SD values are indicated in grey error bars. The phragmoplast microtubules 3 min and 15 min after anaphase onset were analysed in (f) and (g), respectively: (f), control, n = 6 cells; *kin12-IIabcd-2*, n = 6 cells. (g), control, n = 5 cells; *kin12-IIabcd-2*, n = 6 cells. (h) Snapshots of a dividing protonemal tip cell expressing mCherry–α-tubulin (magenta) and MAP65c–citrine (green). The images were captured 3 min after anaphase onset. Scale bar, 10 µm. (i) Normalised intensity profile of mCherry–α-tubulin (magenta) and MAP65c–citrine (green) in the phragmoplast of the control line (left) and the *kin12-IIabcd-1* mutant (right). The intensity was measured perpendicular to the division plane. The midzone position marked by MAP65c–citrine was plotted at 0 µm in the graphs. The mean intensity is depicted with a line, and bars indicate SD values. Control, n = 12 cells; *kin12-IIabcd-1*, n = 9 cells.

### Kinesin12-II facilitates cell plate formation by promoting cell plate material accumulation

Phragmoplast disassembly requires cell plate formation; chemical inhibition of cell plate formation by brefeldin A (BFA), an inhibitor of membrane trafficking from the ER to Golgi apparatus, perturbs phragmoplast disassembly in tobacco BY-2 cells ^48^. To test if this is the case in moss protonemal cells, we treated cells expressing SCAMP4–mNG and mCherry–α-tubulin with BFA. Upon BFA addition, SCAMP4–mNG signals initially accumulated at the midzone but dispersed after 135 min (Supplementary Fig. 5a–c and Supplementary Video 5). The microtubule gap at the midzone was not detected, indicating that the phragmoplast was not promptly disassembled in the presence of BFA (Supplementary Fig. 5d, e). These results suggested that BFA treatment inhibits cell plate formation by perturbing the membrane remodelling process, leading to the inhibition of phragmoplast disassembly in the moss protonemal cells. Thus, the phragmoplast disassembly phenotype observed in the *kin12-IIabcd-2* mutant may be attributed to defects in cell plate formation.

To examine whether cell plate formation is affected by *Kinesin12-II* disruption, we introduced the SCAMP4–mNG marker into the *kin12-IIabcd-1* mutant. Strikingly, we observed that the timing of the appearance of the linear SCAMP4–mNG signal at the phragmoplast midzone was significantly delayed in the *kin12-IIabcd-1* mutant (∼5 times longer after anaphase onset than in the control) (Fig. 4a, b and Supplementary Video 6). A delay was also observed for another cell plate-localised transmembrane proteins, the callose synthase CalS5 ^49,50^ (Supplementary Fig. 6a, b and Supplementary Video 7). These results strongly suggested that cell plate formation was delayed in the mutant. Ectopic expression of Kinesin12-IIc restored the timing of SCAMP4 accumulation and phragmoplast disassembly (Fig. 4b, c), confirming that cytokinesis failure accompanied by delayed cell plate formation was due to the disruption of *Kinesin12-II* genes.

**Figure 4.**
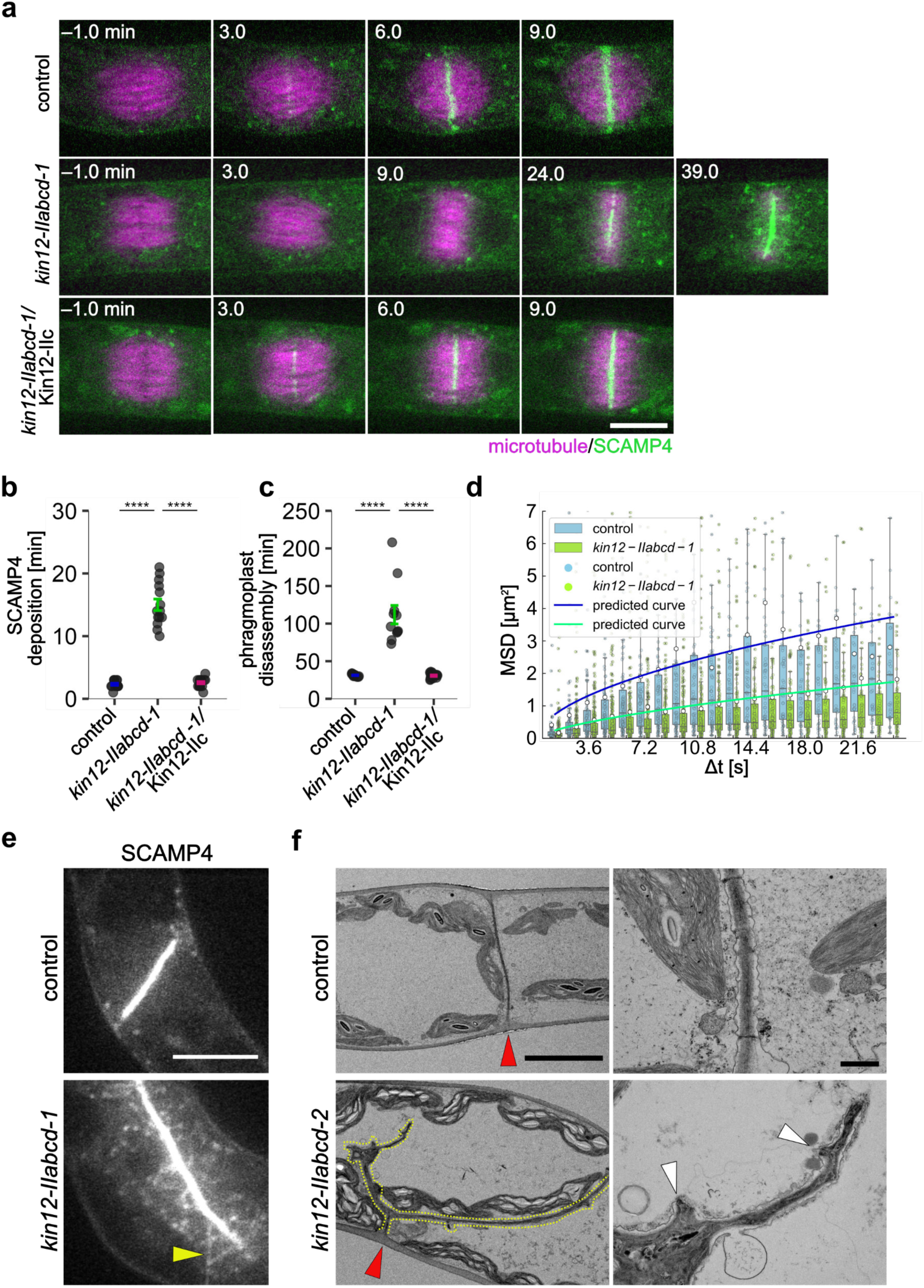
Kinesin12-II is required for cell plate material transport and proper cross-wall formation. (a) Time-lapse images of a dividing protonemal tip cell expressing mCherry–α-tubulin (magenta) and SCAMP4–mNG (green). A single plane of confocal images is presented. In the *kin12-IIabcd-1*/Kin12-IIc moss, the Kin12-IIc protein was ectopically expressed under its native promoter. Time 0 corresponds to anaphase onset. Chloroplast autofluorescence is also faintly visualised in the green channel. Scale bar, 10 µm. (b, c) Comparison of mitotic duration and phragmoplast disassembly duration. (b) Control, 2.3 ± 0.2 min (mean ± SEM), n = 13; *kin12-IIabcd-1*, 15.0 ± 0.9 min, n = 14; *kin12-IIabcd-1*/kin12-IIc, 2.6 ± 0.2 min, n = 16. *P*-value was calculated using ART ANOVA and Tukey’s test, *P* < 0.0001 (control– *kin12-IIabcd-1*), *P* < 0.0001 (*kin12-IIabcd-1*– *kin12-IIabcd-1*/Kin12-IIc); (c) control, 31.2 ± 0.5 min (mean ± SEM), n = 12; *kin12-IIabcd-1*, 111.8 ± 12.4 min, n = 11; *kin12-IIabcd-1*/Kin12-IIc, 30.7 ± 1.0 min, n = 13. *P*-value was calculated using ART ANOVA and Tukey’s test, *P* < 0.0001 (control– *kin12-IIabcd-1*), *P* < 0.0001 (*kin12-IIabcd-1*– *kin12-IIabcd-1*/Kin12-IIc). (d) MSD of SCAMP4 signals in the phragmoplast, plotted against lag time. Analysed images were acquired at 1.2 s intervals. Box plots show median (horizontal lines) and mean (white dots) values; error bars show SD values. *P*-values calculated by bootstrap-based resampling and adjusted by the Holm method were <0.05 at every time point. Curve fittings are depicted as solid blue (control) and green (*kin12-IIabcd-1*) lines. Control; n = 35–114 trajectories, *kin12-IIabcd-1*; n = 76–313 trajectories. (e) Images of a dividing protonemal tip cell expressing SCAMP4–mNG. A single plane of confocal images is presented. The arrowhead indicates a filamentous structure composed of SCAMP4–mNG in the phragmoplast. Scale bar, 10 µm. (f) Transmission electron micrographs of cross-walls in the control and the *kin12-IIabcd-2* mutant. Red arrowheads indicate cross-wall positions, and yellow dashed-lines highlight branched cross-walls in the *kin12-IIabcd-2* mutant. Scale bars, 10 µm (left) and 1 µm (right, magnified images).

### Kinesin12-II is required for directional transport of cell plate materials

Tracking individual SCAMP4–mNG puncta in the phragmoplast revealed that they moved diffusively, rather than directionally, in the *kin12-IIabcd-1* mutant (Supplementary Video 8). We calculated the mean squared displacement (MSD) of individual signals in the presence and absence of the Kinesin12-II protein; the MSD in the control was significantly larger than that in the *kin12-IIabcd-1* mutant (Fig. 4d). In addition, we fitted the calculated MSD as a function of the time lag to the formula:

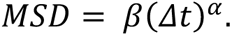

The range of motion represented by the coefficient β in the *kin12-IIabcd-1* mutant was estimated to be three-fold smaller than that of the control (control, α = 0.54 ± 0.11, β = 0.67 ± 0.19; *kin12-IIabcd-*1, α = 0.65 ± 0.07, β = 0.22 ± 0.04). These results indicated that the presence of Kinesin12-II increases directional movement and suggested that Kinesin12-II is involved in the transport of cell plate materials in the phragmoplast.

In addition to the abnormal dynamics, we observed filamentous signals of SCAMP4–mNG that branched out from the midzone signal in the *kin12-IIabcd-1* mutant (Fig. 4e, arrowhead). This was not observed in the control cells. The filamentous structures began to form during phragmoplast expansion and did not disassemble after minutes of observation, suggesting that the cell plate material had ectopically fused and solidified outside the midzone in the phragmoplast.

Next, we examined the protonemal tissue using transmission electron microscopy (TEM). TEM observations revealed that the cross-wall was abnormally formed in the *kin12-IIabcd-2* mutant; the thickness and density of the wall were not uniform, and protrusions were observed, consistent with the mistargeting of the cell plate material in the phragmoplast (Fig. 4f). Thus, Kinesin12-II-mediated rapid and targeted accumulation of cell plate materials may be important for the proper formation of the cross-wall.

### Kinesin12-II exhibits processive motility along the microtubule *in vitro* and *in vivo*

To examine the motor activity of Kinesin12-II on microtubules, we performed two *in vitro* assays using a recombinant Kinesin12IIc^motor^–superfolder (sf)GFP protein (129–523 a.a.) (Fig. 5a, b). First, a microtubule gliding assay was conducted, in which motors were immobilised on a coverslip and then GMPCPP-stabilised microtubules were supplied. Second, the purified Kinesin12IIc^motor^–sfGFP was subjected to a single-motor motility assay, in which microtubules were attached to a coverslip and then a small amount of the motor was supplied. Kinesin12IIc^motor^–sfGFP showed robust microtubule-gliding activity and processive motility in these assays (Fig. 5c–g and Supplementary Video 9). The mean velocities were 120 ± 30 (mean ± SD, n = 182) and 80 ± 70 nm/s (mean ± SD, n = 223), respectively. However, in the motility assay, we did not observe extended dwells at the microtubule ends when the moving motor reached the microtubule end. In addition to directed motility, we observed the diffusive motion of Kinesin12IIc^motor^–sfGFP on the microtubules (Fig. 5e). This property is consistent with that of *Arabidopsis* Kinesin12-I (Kin12D/POK2) ^51^; mode switching might be a conserved feature of plant Kinesin-12. From these *in vitro* analyses, we concluded that Kinesin12-IIc is a motor with dual modes: one processive and the other diffusive along the microtubule.

**Figure 5.**
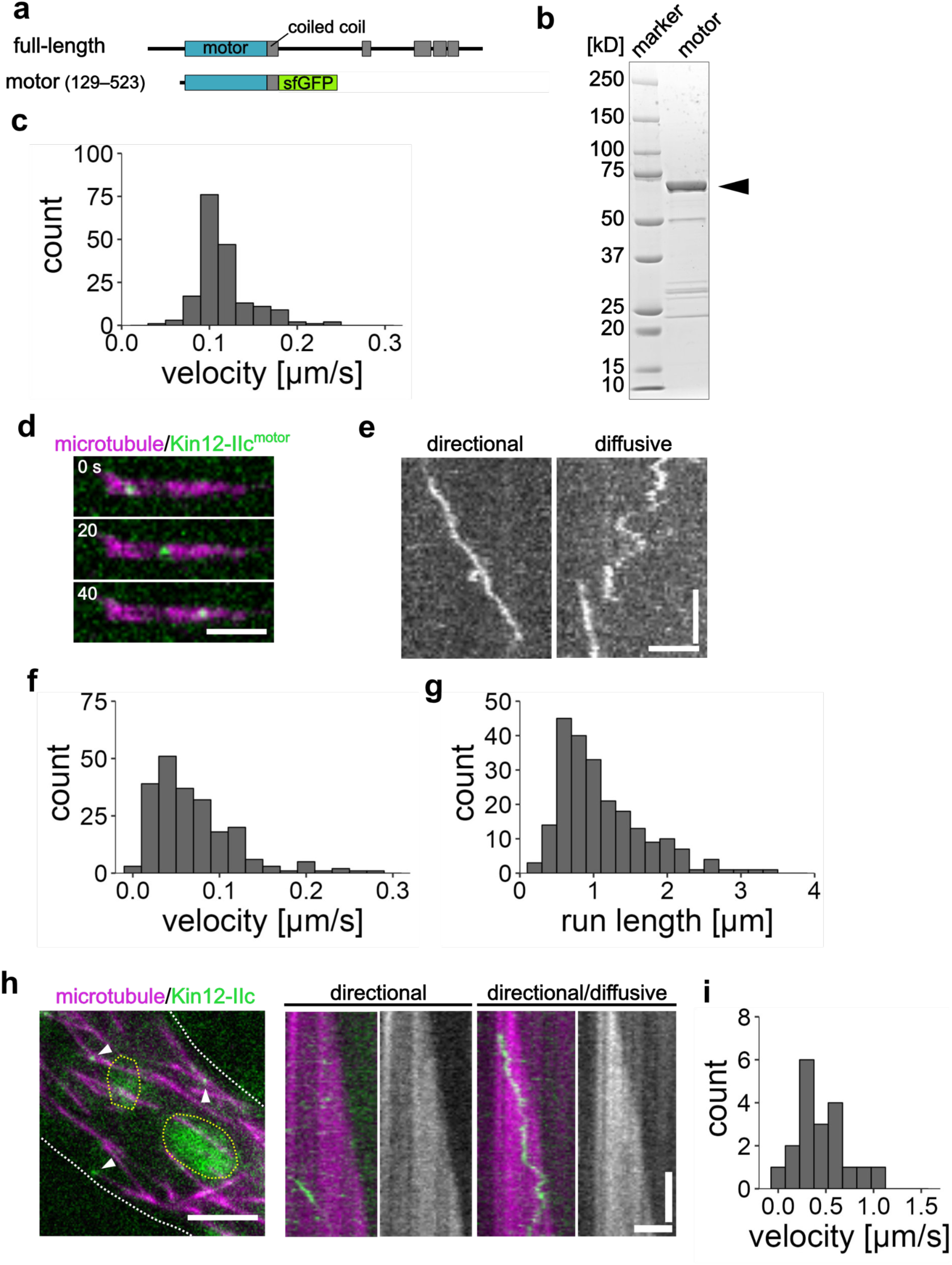
Kinesin12-II moves processively along the microtubule *in vitro* and *in vivo*. (a) Diagrammatic representation of Kinesin12-II construct used for *in vitro* experiments. (b) Coomassie-stained gel of purified Kinesin12-IIc (129–523 a.a.)–sfGFP (Kinesin12-IIc^motor^–sfGFP) protein. The arrowhead indicates a predominant band of approximately the expected size (71.8 kD). (c) Quantification of gliding velocity. 0.12 ± 0.03 µm/s (mean ± SD, n = 182) (d) Sequential images from a single-molecule motility assay. Microtubules (magenta) and Kinesin12-IIc^motor^–sfGFP (green) are presented. Arrowheads indicate the positions of Kinesin12-IIc^motor^–sfGFP moving processively along the microtubule. Scale bar, 2 µm. (e) Kymographs depicting processive and diffusive motility of Kinesin12-IIc^motor^–sfGFP. Horizontal bar, 2 µm; vertical bar, 20 s. (f, g) Quantification of velocity and run length measured in the motility assay. (f) Velocity: 0.08 ± 0.07 µm/s (mean ± SD, n = 223); (g) run length: 1.13 ± 0.76 µm (mean ± SD, n = 223). (h) Image of a protonemal cell expressing mCherry–α-tubulin (magenta) and Kin12-IIc–mNG (green), acquired with oblique TIRF microscopy and kymographs showing directional and diffusive movements of Kin12-IIc–mNG. Arrowheads indicate Kin12-IIc-mNG associated with the microtubule. Cell edges are marked by white dashed lines, and chloroplast autofluorescence is encircled with yellow dashed lines. Scale bar (left), 5 µm; horizontal bar (right), 2 µm; vertical bar (right), 5 s. (i) Quantification of Kin12-IIc–mNG velocity *in vivo*—0.46 ± 0.27 µm/s (mean ± SD, n = 19).

Next, we visualised the motility of individual Kinesin12-II proteins in the phragmoplast. However, it was impossible to observe them because the faint signals along the phragmoplast microtubules were masked by extremely strong fluorescent signals at the midzone. Therefore, we instead observed the endoplasm of the protonemal cell using oblique illumination microscopy ^52^, with which we were able to detect a small amount of Kinesin12-II occasionally associated with the microtubule (Fig. 5h left). We observed that Kinesin12-IIc–mNG signals moved processively as well as diffusively along the endoplasmic microtubule (processive motility: 460 ± 270 nm/s, mean ± SD, n = 19; Fig. 5h right, 5i and Supplementary Video 10). The processive movement was plus-end-directed, as Kinesin12-IIc–mNG moved towards the direction of persistent microtubule growth (Fig. 5h). Based on these observations, we concluded that Kinesin12-IIc is a plus-end-directed motor capable of carrying cargo.

### Non-motor regions are required for midzone accumulation

Members of the Kinesin12-II subfamily including *Arabidopsis* Kinesin12-II possess a motor domain at the N-terminus, followed by clusters of coiled coils in the tail region (Fig. 6a) ^29^. We functionally dissected the Kinesin12-IIc protein by transforming truncated or mutated constructs tagged with mNG into the *kin12-IIabcd-1* mutant (Fig. 6a). After transgenic line selection, we assessed whether the growth phenotypes were restored by the ectopic expression of the constructs. Among seven constructs we transformed, we could not obtain stable transgenic lines for the non-motile “rigor” (T227N) and headless (473–1345 a.a.) mutants, probably due to the dominant negative effect of these constructs^53–55^. Transgenic lines were successfully obtained for five other constructs. However, none of the truncated constructs complemented plant growth retardation or gametophore developmental disorder (Fig. 6a, b and Supplementary Video 11). The “motor” construct, which contains only the motor domain and the first coiled coil for protein dimerization, decorated the entire phragmoplast microtubules without enrichment at the midzone. Considering that Kinesin12-IIc^motor^–sfGFP exhibited processive motility *in vitro* and Kinesin12-IIc–mNG did not show an extended dwell at the microtubule end (Fig. 5e, h), this result suggests that processive motility is insufficient for midzone localisation. The “delta C” and “middle” constructs were localised at the midzone, despite the lack of a motor domain in the latter. In contrast, the “C-terminus” construct was not detected in the phragmoplast in the early cytokinetic phase in the *kin12-IIabcd-1* mutant (Fig. 6a). It exhibited a punctate signal and faint accumulation in the phragmoplast during late cytokinesis in the mutant, at which point the cell plate was partially formed (Fig. 6c and Supplementary Video 12). However, when expressed in the control line, the same construct was enriched at the midzone from the early cytokinetic phase (Fig. 6c). We interpreted that this region interacts with the cell plate rather than phragmoplast microtubules. The “C-terminus” construct was also detected at the cell apex, where actin filaments, microtubules and exocytic vesicles are enriched to propel tip growth ^56^ (Supplementary Fig. 7). The localisation pattern of the C-terminus-truncated proteins at the apex was similar to the vesicle distribution pattern, rather than to that of microtubules or actin ^57^. These results suggest that Kinesin12-II may interact via the C-terminal region with common vesicular proteins that localise at the cell plate and apex. Finally, we examined whether a truncated protein containing a motor domain and putative cell plate recognition motif could restore the phenotype. To this end, we expressed the “delta middle” construct in the mutant. We observed that it was localised at the midzone at lower levels than the full-length or middle constructs, and the growth phenotype was not rescued (Fig. 6a). These results indicated that the middle region is required for stable midzone localisation of kinesin-12-IIc and for cell plate formation.

**Figure 6.**
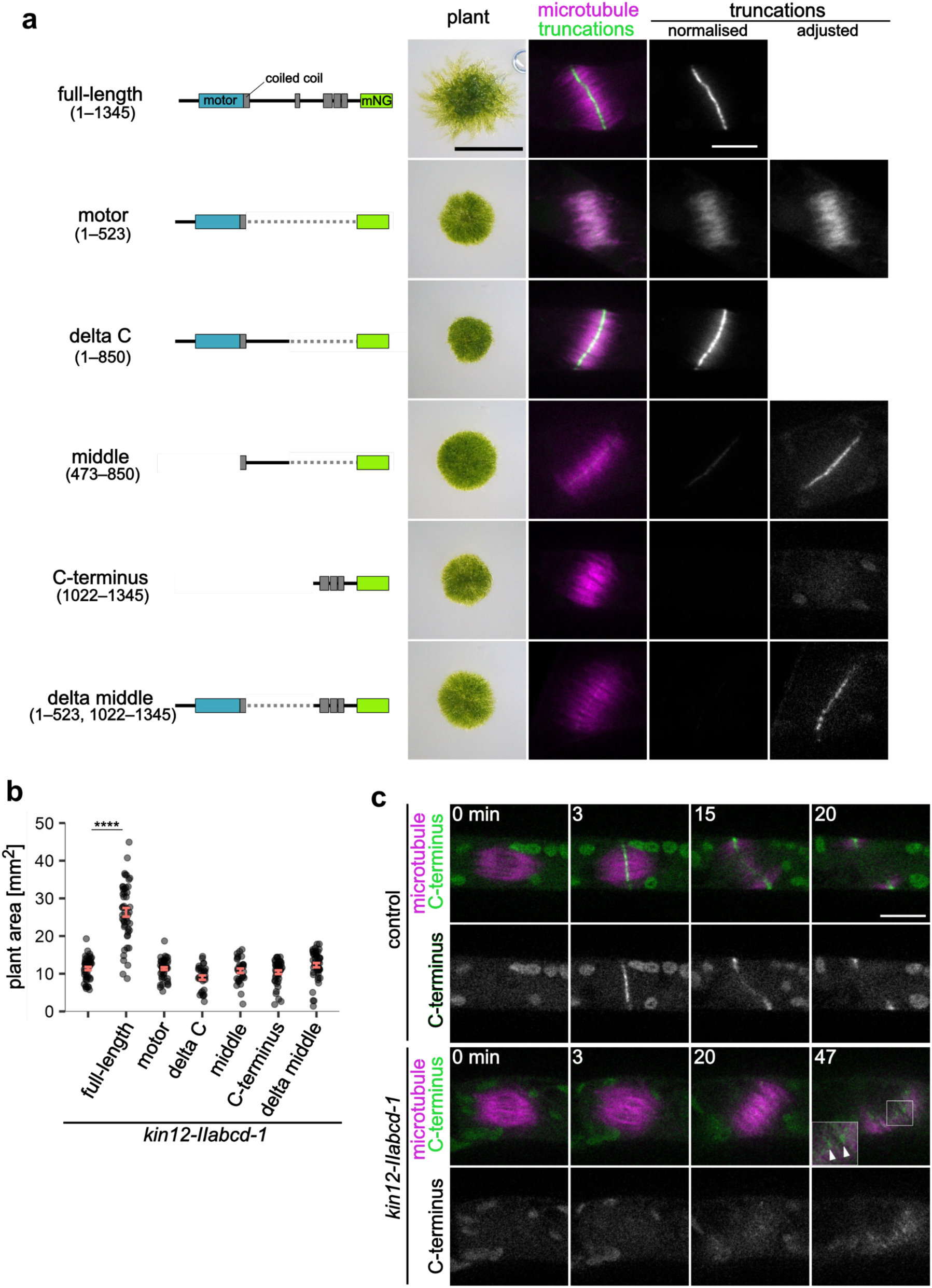
Non-motor regions are required for midzone accumulation. (a) Schematic representation of truncated Kinesin12-IIc constructs used in subsequent analysis and images showcasing 3-week-old moss and time-lapse snapshots of a dividing protonemal cell expressing mCherry–α-tubulin (magenta) and truncated Kin12-IIc–mNG variants (green). In the “normalised” and “adjusted” columns, intensities of the mNG signals were normalised and individually adjusted, respectively. The phragmoplast images were acquired 15 min after anaphase onset. Chloroplast autofluorescence is also faintly visualised in the green channel. Scale bar (left), 5 mm; scale bar (right),10 µm. (b) Analysis of the moss growth area over three weeks, cultured from protoplasts; *kin12-IIabcd-1*, 11.4 ± 0.4 mm^2^ (mean ± SEM), n = 48; *kin12-IIabcd-1*/full-length, 26.3 ± 1.1 mm^2^, n = 48; *kin12-IIabcd-1*/motor, 11.4 ± 0.4 mm^2^, n = 48; *kin12-IIabcd-1*/delta C, 9.0 ± 0.6 mm^2^, n = 30; *kin12-IIabcd-1*/middle, 10.8 ± 0.6 mm^2^, n = 31; *kin12-IIabcd-1*/C-terminus, 10.4 ± 0.5 mm^2^, n = 44; *kin12-IIabcd-1*/delta middle, 12.3 ± 0.6 mm^2^, n = 46. *P*-value was calculated using ART ANOVA and Tukey’s test; *P* < 0.0001 (*kin12-IIabcd-1*– *kin12-IIabcd-1*/full-length) and *P* > 0.05 (*kin12-IIabcd-1*– *kin12-IIabcd-1*/motor, *kin12-IIabcd-1*– *kin12-IIabcd-1*/delta C, *kin12-IIabcd-1*– *kin12-IIabcd-1*/middle, *kin12-IIabcd-1*– *kin12-IIabcd-1*/C-terminus, and *kin12-IIabcd-1*– *kin12-IIabcd-1*/delta middle). (c) Time-lapse snapshots of a dividing protonemal cell expressing mCherry–α-tubulin (magenta) and Kin12-IIc (C-terminus)–mNG (green). Time 0 corresponds to anaphase onset. Chloroplast autofluorescence is also faintly visualised in the green channel. Scale bar, 10 µm.

### *Kinesin12-II* disruption causes chromosome missegregation in the early gametophore

In contrast to the unique protonemal tissue, gametophores share many features with multiple organs in flowering plants, such as three-dimensional development (Fig. 7a)^58^. The early gametophore repeats asymmetric cell division to maintain stem cells at the apex. We found that the cell cycle duration ranged from 5 to 8 h in gametophore initial cells (Fig. 7b), which is equivalent to that of the apical cells of the caulonemal filaments, but much shorter than that of the apical cells of the chloronemal filaments (22–26 h) ^59^. Since the characteristic phenotype caused by *Kinesin12-II* disruption involved a lack of mature gametophores (Fig. 2), we hypothesised that the gametophore’s developmental process was impeded by *Kinesin12-II* disruption. To address the mechanism of gametophore developmental disorders, we performed time-lapse imaging of early gametophores. Consistent with the phenotype observed in protonemal cells, phragmoplast disassembly was delayed in the *kin12-IIabcd-2* mutant and phragmoplast microtubules were disorganised during prolonged cytokinesis (Fig. 7c and Supplementary Video 13). Strikingly, cytokinesis often took longer than the entire cell cycle of gametophore initial cells of the control line (Fig. 7d). Consequently, the mutant cells entered the mitotic phase of the subsequent cell cycle before the complete disassembly of the phragmoplast in the previous round, and the remaining phragmoplast microtubules were integrated into the spindle, causing chromosome missegregation and the production of aneuploid cells (Fig. 7e, f). Aneuploidy is a common cause of growth and developmental disorders ^60^, which explains the halt in gametophore development in the *kinesin12-II* mutants. These results illustrated that the rapid completion of cell plate formation is indispensable for organ development.

**Figure 7.**
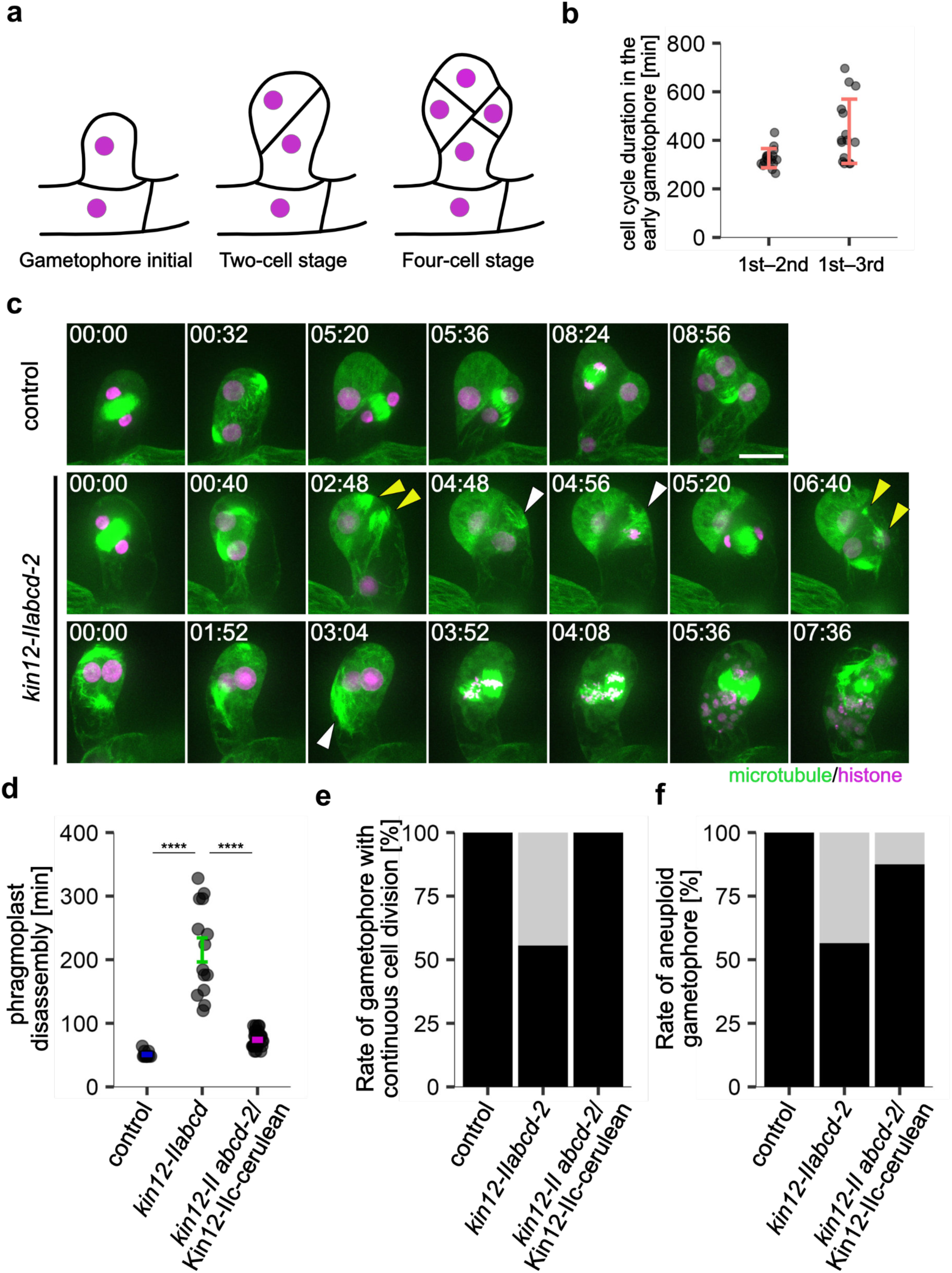
Kinesin12-II is required for proper chromosome segregation in the early gametophore. (a) Schematic representation of the early stages of gametophore development bulged from the sub-apical cell in protonemal cells. Circles represent nuclei. (b) Measurement of the cell cycle duration in early gametophores. Duration from NEBD of the first division to NEBD of the second or third division were measured. Error bars represent SD values. Duration from 1^st^ to 2^nd^ division; 326 ± 39 min (mean ± SD), n = 16; 1^st^ to 3^rd^ division; 437 ± 132 min, n = 15. (c) Sequential images of a dividing gametophore initial cell expressing GFP–α-tubulin (green) and histoneH2B–mCherry (magenta). Maxman intensity projection images of 21 confocal images, acquired every 2 µm, are presented. The value 00:00 represents 0 h 0 min. Scale bar, 20 µm. (d) Comparison of phragmoplast disassembly duration. Control, 50.9 ± 1.7 min (mean ± SEM), n = 11; *kin12-IIabcd-2*, 215.4 ± 18.9 min, n = 14; *kin12-IIabcd-2*/Kin12-IIc– cerulean, 74.0 ± 2.2 min, n = 32. *P* value was calculated using ART ANOVA and Tukey’s test, *P* < 0.0001 (control– *kin12-IIabcd-2*), *P* < 0.0001 (*kin12-IIabcd-2*– *kin12-IIabcd-2*/Kin12-IIc). (e) Proportion of gametophores exhibiting continuous cell division (cells entering the next division before completion of phragmoplast disassembly). Normal cell division is represented in black and continuous cell division in grey. Control, n = 16; *kin12-IIabcd-2*, n = 18; *kin12-IIabcd-2*/Kin12-IIc-cerulean, n = 32 (f) Proportion of gametophores with aneuploid cells. Normal gametophore in black, gametophore with aneuploid cells in grey. Control, n = 16; *kin12-IIabcd-2*, n = 23; *kin12-IIabcd-2*/Kin12-IIc-cerulean, n = 32.

## Discussion

In the present study, we addressed a long-standing question concerning the mechanisms by which cell plate materials are targeted and accumulate at the phragmoplast midzone. Our series of live-imaging analyses supports a model wherein Kinesin12-II is responsible for transporting vesicles along the phragmoplast microtubules and retaining them at the midzone, thus facilitating efficient cell plate formation during cytokinesis. Our data suggest that this motor protein-mediated cell plate formation system is indispensable for multicellular organ development.

### Moss Kinesin12-II functions not only as a transporter but also as a retention factor of cell plate materials in the phragmoplast

Electron microscopic observations have assumed the presence of motor-dependent vesicular transport in plant cells ^24^. However, empirical evidence of this concept has been lacking. Using the moss protonemal tissue, we observed the directional movement of cell plate materials marked by SCAMP4–mNG. Furthermore, this movement was abolished by *Kinesin12-II* disruption. The Kinesin12-II motor protein showed plus-end-directed processive motility along the microtubule*s in vitro* and *in vivo*. Consequently, we propose that moss Kinesin12-II is a long-assumed motor that acts as a transporter of cell plate materials towards the midzone in the phragmoplast.

In this study, we used the SCAMP4 protein as a vesicular marker. SCAMP proteins have been used as markers for secretory vesicles originating from the trans-Golgi network in plant cells ^35,61^. In *P. patens*, SCAMP4 localises to early endosomal compartments, the plasma membrane and the cell plate ^10^. The SCAMP4-marked punctate signals largely overlap with those of the lipophilic dye FM4-64, suggesting that various types of vesicles are marked with SCAMP4 in the moss ^10^. However, it cannot be ruled out that Kinesin12-II is responsible for the transport of SCAMP4-free vesicles, which might include CalS5-containing vesicles, whose midzone accumulation was also significantly delayed in the *kin12-IIabcd-1* mutants. Kinesins typically bind to cargo vesicles via specific adaptor or scaffold proteins, thereby enabling directional transport towards specific targets ^25^. Given that our observations suggest that the C-terminus of Kinesin12-II interacts with the cell plate rather than the phragmoplast microtubules, it is plausible that Kinesin12-II recognises its cargo through the C-terminus. Identifying the direct interacting partner of the C-terminus will be a key future investigation in comprehending the versatility of Kinesin12-II cargos.

The phragmoplast midzone, where the antiparallel microtubule overlap is formed, plays a pivotal role in spatially confining vesicle immobilisation and facilitating localised fusion ^10^. While the precise mechanism by which cell plate vesicles associate with this cytoskeletal structure remains unclear, our data imply the potential involvement of Kinesin12-II in the retention of vesicles at the midzone. Neither *in vitro* experiments utilising a processive Kinesin12-II motor construct nor time-lapse observations of Kinesin12-IIc in endoplasmic microtubules showed extended dwells at microtubule ends. This suggests that Kinesin12-II on its own dissociates from the microtubule upon reaching the plus-end within the midzone, which is different from other kinesins, such as kinesin-8, that have long dwell times at the plus-end ^62–64^. Nevertheless, Kinesin12-II does accumulate at the midzone, indicating that Kinesin12-II is retained by a motor-independent mechanism. Domain dissection analysis provided insights into the localisation mechanism. Specifically, the middle region and delta C construct were preferentially localised at the midzone, whereas the motor domain was found along the entire phragmoplast microtubule without midzone enrichment. This suggests that the middle region is responsible for the retention of Kinesin12-II at the midzone. Moreover, the delta middle construct comprising the motor and C-terminal regions exhibited weak localisation at the midzone and was insufficient to restore the phenotype, suggesting the importance of the middle region for stable localisation. The question arises as to why stable localisation of Kinesin12-II at the midzone is required, especially considering that the midzone is the terminus for cell plate material transport. A plausible hypothesis is that Kinesin12-II is entrapped at the midzone after cargo transport, potentially immobilising these cargos, thereby facilitating efficient membrane remodelling at the CPAM. This corroborates a previous observation that bipolar phragmoplast microtubules lacking antiparallel microtubule interdigitation in cells depleted of MAP65 are insufficient for the sustained accumulation of cell plate-destined vesicles ^13^. We propose that Kinesin12-II acts not only as a transporter but also as a retention factor for the cell plate precursor at the midzone.

### Why do plant cells utilise the energy-consuming transport mechanism over diffusion for cytokinetic vesicle accumulation?

Generally, motor-driven active transport along the cytoskeletal filament can carry cargo directly to a target, thereby surpassing the speed of passive diffusion. In the *kin12-IIabcd-1* mutant, which lacks directional vesicle transport in the phragmoplast, the MSD was significantly lower than that in the control. This implies that the transport mechanism facilitates the delivery of cell plate materials, resulting in rapid formation of the cell plate. Observations of the cross-wall and time-lapse imaging of the early gametophore in the *kinesin12-II* mutants underscore the necessity for rapid accumulation and formation of the cell plate, respectively. In the mutant lacking Kinesin12-IIs, SCAMP4 deposition at the midzone was significantly delayed, and the SCAMP4 signal branched out from the midzone in the phragmoplast. TEM revealed that the mutant formed branched and disordered cross-walls. This suggests that membrane remodelling can occur ectopically in phragmoplasts in the absence of material transport, emphasising the need for rapid and targeted material accumulation prior to membrane remodelling. Moreover, the early gametophore cell with the expanding phragmoplast entered subsequent cell division, resulting in arrested cell plate formation in the absence of Kinesin12-II. Simultaneously, the remaining phragmoplast microtubules were integrated into the spindle formed for subsequent cell division. This sequence of events triggered chromosome missegregation and aneuploidy, and halted gametophore development. Consequently, it is inferred that moss cells employ a Kinesin12-II-mediated vesicle transport system to establish a uniform cross-wall and facilitate the sequential cell division cycle during multicellular organ development.

### Is Kinesin12-II’s function conserved in other plants?

The conservation of Kinesin12-II function across various plant species presents an intriguing question. A previous study regarding AtKin12A/AtKin12B function was conducted using *Arabidopsis* microspores. Kin12A and Kin12B have been suggested to redundantly organise phragmoplast microtubules into two mirrored sets with a gap between them by capturing the plus-ends of the microtubules. The loss of microtubule gap formation at the midzone is presumed to be the primary cause of cell plate formation defects in the *Arabidopsis kin12a kin12b* double mutant ^33^. In contrast, our findings demonstrate the proper organisation of phragmoplast microtubules into two opposing sets in moss protonemal cells, even upon *Kinesin12-II* disruption: MAP65c localised at the midzone, and the majority of EB1 signals moved towards the midzone in the absence of Kinesin12-II. It is possible that moss Kinesin12-IIs have acquired functions distinct from those of Kin12A and Kin12B in *Arabidopsis*. However, the characteristic phenotypes observed in the *Arabidopsis* mutant, such as the absence of gap formation at the midzone, the collapse of the phragmoplast and the lack of cell plate material accumulation, were also observed in the moss mutant. Previous results have suggested that Kin12A and Kin12B depletion diminishes cell plate material transport, leading to the loss of microtubule gap formation. This scenario is supported by the BFA experiment, in which the inhibition of cell plate formation precluded microtubule gap formation. It will be interesting to revisit this phenotype and examine how cell plate material transport is affected in the *kin12a kin12b* mutant.

Kinesin12-II, despite its homological classification in the Kinesin-12 family alongside animal KIF15, has evolved plant-specific roles, particularly in cell plate material transport. Recent phylogenetic analyses of motor proteins among Chlorophyta and Streptophyta have suggested that Kinesin12-II is present in *Chara braunii* and land plants, but not in Chlorophyta ^65^. The emergence of Kinesin12-II intriguingly coincides with the development of the phragmoplast-mediated cell plate formation system during plant evolution; the ancestral cytokinesis system in Streprophyta involved centripetal cleavage, whereas the phragmoplast system arose prior to the evolutionary branching of Charophyceae ^66^. Therefore, it is conceivable that plants have diversified the functions of *Kinesin-12* genes in line with the evolution of a unique cell division system.

## Materials & Methods

### Molecular cloning and gene targeting experiments

All transgenic *Physcomitrium patens* lines, plasmids, primers and amino acid sequences of the mutants used in this study are listed in Supplementary Tables 1, 2, 3 and 4. These moss lines originate from the Gransden ecotype of *Physcomitrium patens*. Moss culture, transformation and selection of transgenic lines were performed in accordance with previously established protocols ^67^. In brief, moss cells were routinely cultured on BCDAT medium under conditions of continuous light illumination at 23–25°C. The transformation procedure was a standard PEG-mediated method.

A series of *kinesin12-II* mutants were generated in the moss strain expressing either GFP-α– tubulin/histone-H2B–mCherry or mCherry–α-tubulin using CRISPR/Cas9-mediated gene editing^13,68,69^. CRISPR targeting sites were manually selected around the characteristic SSRSH motif (Switch 1 element) and the LAGSE motif (Switch 2 element) within the motor domain, both of which are indispensable for ATP hydrolysis, employing the CRISPOR program ^70^. Target sequences were synthesised and integrated into the BsaI site of pPY157, a vector comprising a U6 promoter and a RNA scaffold. Subsequently, multiple CRISPR cassettes were cloned into a vector carrying nourseothricin resistance using the In-Fusion HD Cloning Kit according to methods described previously ^46^. For the transformation, a mixture comprising 5 µg of the circular multicassette plasmid, 30 µg of *Streptococcus pyogenes* Cas9 expression vector (pGenius) and 10 µl of 50 µM synthesised oligonucleotide templates was applied to the moss^46,71^. Finally, the mutations were confirmed by genotyping PCR, followed by sequencing.

For endogenous tagging via homologous recombination, the plasmid was constructed using an In-Fusion HD Cloning Kit. Approximately 1 kilobase (kb) sequences both upstream and downstream of the stop codon of the genes of interest flanking an in-frame linker, fluorescent protein-coding sequence, Flag tag and an antibiotic resistance cassette constituted the homologous recombination template for C-terminal tagging. For N-terminal tagging, 1 kb sequences both upstream and downstream of the start codon of the genes of interest were utilised. The transgenic line was confirmed by PCR using primers specifically designed to target the regions upstream and downstream of the homologous recombination template of the genes of interest.

For ectopic expression plasmids, the coding sequence was amplified from the moss complementary DNA library and integrated into a vector containing a promoter sequence, in-frame linker (GSGGSG), mNG-coding sequence, Flag tag, hygromycin resistance cassette and 1 kb sequences homologous to the *hb7* locus. In these ectopic expression experiments, either the *P. patens* EF1α promoter sequence or a 2 kb upstream sequence from the start codon of the *Kinesin12-IIc* gene was employed to drive protein expression. The transgenic line was confirmed by either PCR or observation of fluorescent protein expression.

### Moss growth assay

The protoplast regeneration assay was performed following a previously described method with some modifications ^67^. Briefly, the moss digested on cellophane-lined BCDAT plates was treated with an 8% mannitol solution supplemented with 0.8% driselase. The resulting protoplasts were washed with an 8% mannitol solution and resuspended in 7 ml of protoplast regeneration liquid. Following an overnight incubation in the dark, the protoplasts were centrifuged at 510 × *g* for 2 min and resuspended in PRM/T agar. The protoplast solution was then spread onto cellophane-lined PRM plates. Subsequently, the protoplasts were cultured for 3 d and then transferred to a BCDAT plate, followed by a 4-day culture. Seven-day-old protonemal cells were imaged using a stereomicroscope SMZ800N (Nikon) equipped with an ILCE-QX1camera (SONY). Ten-day-old mosses were individually transferred to BCDAT plates and cultured for 3–5 weeks. Images of whole mosses and gametophores were captured using a COOLPIX3100 digital camera (Nikon, Tokyo, Japan) and SMZ800N, respectively.

### Imaging conditions

Spinning disc confocal microscopy was performed using a Nikon Ti inverted microscope equipped with lenses of varying magnifications:100×1.45-NA lens, 60×1.40-NA lens and 40×1.30-NA lens. Additional equipment included an ImagEM EMCCD camera (Hamamatsu Photonics) and a CSU-X1 spinning-disc confocal scanner unit (Yokogawa), utilised as previously described (Yamada et al., 2016). Z-series images were captured using Nano-Z Series nanopositioner combined with a Nano-Drive controller, with z-stack imaging executed at 1–2 µm intervals. Lens selection was based on the specific requirements of each experiment. Oblique illumination fluorescence microscopy and *in vitro* microscopy were conducted using a Nikon Ti inverted microscope with a TIRF unit, a 100× 1.49-NA lens, a GEMINI split view (Hamamatsu Photonics) and an Evolve EMCCD camera (Roper). The microscope was operated using a NIS-Elements (Nikon). Imaging was performed at room temperature in the dark, except during early gametophore observation, when a light cycle of 4 min illumination and 4 min dark was applied.

### Imaging sample preparation

For confocal time-lapse imaging, moss samples were prepared on an agar pad in 35 mm glass-bottomed dishes, as previously described (Yamada et al., 2016). Briefly, protonema cells were cultured on glass coated with BCD agar for 5–7 d.

To observe early gametophore development, moss agar pad samples were prepared as previously described. One micromolar of 2-isopentenyladenine (2ip), a cytokinin, was added to the 5-day-cultured agar pad sample 22 h before imaging ^72^.

For oblique illumination imaging, moss samples were prepared in a microfluidic device with a 15 µm height as described previously with some modifications ^68^. Briefly, protonemal cells grown on cellophane-lined BCDAT medium for 5–7 d were sonicated and injected into the microfluidic device. The protonemal cells in the device were then immersed in Hoagland Medium and cultured for 3–4 d at room temperature under continuous light.

### Drug treatment

In this study, brefeldin A (BFA, 50 µM) and latrunculin A (5 µM) were employed, with 0.5% dimethyl sulfoxide (DMSO) serving as a control. Stock solutions of BFA (18 mg/ml) and latrunculin A (25 mM) were prepared in DMSO and stored at –20°C. For experimental use, the solutions were diluted to the designated concentrations using water. Drugs were applied to the moss agar pad samples 15–30 min before imaging, as prolonged BFA treatment prevented the cells from entering mitosis.

### Electron microscopy observation

Protonemal cells cultured on cellophane-lined BCDAT medium for six d underwent a two-step fixation process for electron microscopy. Initially, protonemal tissue was immersed in 0.05 M cacodylate buffer (pH 7.4) containing 2% paraformaldehyde and 2% glutaraldehyde. Following overnight incubation at 4°C, the tissue was then washed thrice with 0.05M cacodylate buffer (pH 7.4) for 30 min each.

Subsequent post-fixation involved 0.05 M cacodylate buffer (pH 7.4) containing 2% osmium tetroxide for 3 h at 4°C. The dehydration process entailed a series of ethanol treatments at increasing concentrations: 50% (4°C, 30 min), 70% (4°C, 30 min), 90% (30 min), 99.5% (30 min, thrice) and 99.5% (overnight). Embedding of the tissue was executed through several steps: propylene oxide (30 min, twice), a mixture of propylene oxide and Quetol-651 (Nisshin EM, 3 h), Quetol-651 (3 h) and Quetol-651 (60°C, 48 h). Ultrathin sections (80 nm) were prepared using an Ultracut UCT (Leica) and stained with 2% uranyl acetate for 15 min and Lead stain solution (Sigma) for 3 min. The sections were observed using a JEM-1400Plus (JEOL). All experiments were conducted at room temperature unless otherwise stated.

### Protein purification

The motor domain of Kinesin12-IIc was cloned from the moss cDNA library and integrated into a vector carrying sfGFP and 6×His sequences using In-Fusion (plasmid pMY742). Expression of Kinesin12-IIc ^motor^–sfGFP–6×His was induced in SoluBL21 *E. coli* with 0.2 mM IPTG for 18 h at 18°C. Harvested cells were lysed using the Advanced Digital Sonifier D450 (Branson) in lysis buffer (25 mM MOPS [pH 7.0], 250 mM KCl, 2 mM MgCl2, 1 mM EGTA, 20 mM imidazole, 0.1 mM ATP) supplemented with 5 mM β-mercaptoethanol, 500 U benzonase and protease inhibitors (1 mM PMSF and peptide inhibitor cocktail: 5 mg/mL aprotinin, 5 mg/mL chymostatin, 5 mg/mL leupeptin, 5 mg/mL pepstatin A). After centrifugation, the clarified lysate was incubated with nickel-NTA coated beads for 1.5 h at 4 °C. Following three washes with the lysis buffer, proteins were eluted using 500 µl elution buffer (25 mM MOPS [pH 7.0], 75 mM KCl, 2 mM MgCl2, 1 mM EGTA, 200 mM imidazole, 0.2 mM ATP) supplemented with 5 mM β-mercaptoethanol. The elution was repeated five times and the second and third fractions were combined and aliquoted after the addition of 20% (w/v) sucrose. The aliquots were flash-frozen in liquid nitrogen and stored at –80°C. Protein concentration was determined by Coomassie staining of the purified protein on an SDS–PAGE gel alongside a BSA reference.

### In vitro assay

Microtubule polymerisation was performed by preparing a biotinylated tubulin mixture. This contained 10% biotinylated pig tubulin and 30% Alexa Fluor 568-labeled pig tubulin in a total concentration of 100 µM with 0.5mM GMPCPP, which was prepared in MRB80 (80 mM Pipes-KOH, pH6.8, 1mM EGTA, 4 mM MgCl2). The tubulin mixture was incubated for 35 min at 37°C to induce polymerisation.

For the gliding assay with purified Kinesin12-IIc^motor^–sfGFP, the methods described in a previous study were referenced ^73^, and 1×Standard Assay Buffer (SAB: 25 mM MOPS [pH 7.0], 75 mM KCl, 2 mM MgCl2, 1 mM EGTA) was utilised. Briefly, 7 µL of the purified recombinant protein was introduced into a flow chamber and incubated at room temperature for 2 min in the dark. Subsequently, a 10 µL reaction mixture (1×SAB, 0.1% methylcellulose, 50 mM glucose, 0.5 µg/µL κ-casein, GMPCPP-stabilised microtubule seeds, oxygen scavenger system, 1 mM ATP) was introduced into the flow chamber, and it was sealed using candle wax. TIRF imaging of Alexa Fluor-568 labelled GMPCPP-stabilised microtubule seeds was performed every 3 s for 10 min at 23–25 °C.

For the single-molecule motility assay with purified Kinesin12-IIc^motor^–sfGFP, the methods described in a previous study were referenced with some modifications to the buffer ^74^. A silanised coverslip was coated with an antibiotin antibody, and 1×SAB buffer containing 1% pluronic acid was introduced into the chamber. After washing once with 1×SAB, GMPCPP-stabilised microtubules labelled with Alexa Fluor 568 and biotin were loaded and incubated for 5 min. After washing once with 1× SAB, 10 µL of reaction mixture (1×SAB, 0.1% methylcellulose, 50 mM glucose, 0.5 µg/µL κ-casein, 150 nM Kinesin12-IIc^motor^–sfGFP, 75 mM KCl, oxygen scavenger system, 1 mM ATP) was introduced into the flow chamber, and it was sealed using candle wax. TIRF imaging was performed every 1 s for 10 min at 23–25 °C.

### Data analysis

All raw data were processed and analysed using FIJI software.

#### SCAMP4 signal motility rate

Spinning-disc confocal fluorescence time-lapse imaging was conducted using a 100×1.45-NA lens, capturing images at intervals of 1–1.3 s. Samples from moss protonemal cells expressing SCAMP4– mNG and mCherry–α-tubulin were analysed. Kymographs were generated along the track of the SCAMP4–mNG, and the velocity was calculated from the slopes observed in these kymographs.

#### Microtubule growth rate

Spinning-disc confocal fluorescence time-lapse imaging was conducted using a 100×1.45-NA lens, capturing images at intervals of 3 s. Samples from the imaging of moss protonemal cells expressing EB1b–mNG and mCherry–α-tubulin were analysed. The *mNG* gene was inserted into the C-terminus of the endogenous *EB1b* gene, allowing for the marking of growing microtubule plus-ends by EB1b–mNG at native expression levels. The surface of the phragmoplast was imaged to observe individual EB1 signals. Kymographs were generated along the phragmoplast microtubules, and the growth rate was derived from the slopes of EB1b–mNG movement in these kymographs.

#### Plant size comparison

To compare plant sizes, images of growing mosses, prepared as described above, were captured using a COOLPIX3100 digital camera (Nikon). Images were analysed using FIJI software, where they were first converted to 8-bit images, automatically thresholded and then binarised. The mosses were automatically outlined using the wand tool and the resulting area measurements were obtained.

#### Mitotic duration

Mitotic duration was analysed using images captured at intervals of 0.5–1 min with a spinning-disc confocal microscope (either a 100×1.45-NA lens or a 60×1.40-NA lens). Mitotic duration was defined as the time from NEBD (nuclear envelope breakdown) to anaphase onset.

#### Cell plate deposition time

To determine the duration of cell plate deposition, spinning-disc confocal fluorescence time-lapse imaging was conducted using either a 100×1.45-NA lens or 60×1.40-NA lens, capturing images at intervals of 0.5–1 min. Samples from the imaging of moss protonemal cells expressing either SCAMP4– mNG or mNG–CalS5 were analysed. Duration of cell plate deposition was defined as the time from anaphase onset to the appearance of a linear cell plate structure. Anaphase onset was determined based on spindle elongation.

#### Phragmoplast disassembly time

Spinning-disc confocal fluorescence time-lapse imaging was conducted using either a 100×1.45-NA lens or 60×1.40-NA lens, capturing images at intervals of 0.5–1 min. Samples from the imaging of moss protonemal cells expressing either mCherry–α-tubulin or GFP–α-tubulin were analysed. Disassembly time was defined as the period from the onset of anaphase to the complete disappearance of phragmoplast microtubules.

#### Phragmoplast rotation angle

Spinning-disc confocal fluorescence time-lapse imaging was conducted using either a 100×1.45-NA lens or 60×1.40-NA lens, capturing images at intervals of 0.5–1 min. Samples from the imaging of moss protonemal cells expressing GFP–α-tubulin were analysed. The angles of the phragmoplast at the onset of anaphase and the angles of the cross-wall were measured, and the difference between these angles was calculated.

#### Phragmoplast displacement

Spinning-disc confocal fluorescence time-lapse imaging was conducted using either a 100×1.45-NA lens or 60×1.40-NA lens, capturing images at intervals of 0.5–1 min. Samples from the imaging of moss protonemal cells expressing GFP–α-tubulin were analysed. The distance between the centre of the phragmoplast midzone at anaphase onset and the centre of the cross-wall was measured.

#### Intensity plotting along the phragmoplast

For intensity plotting along the phragmoplast, images were captured every 0.5–1 min using the spinning-disc confocal fluorescence microscope (either 100×1.45-NA lens or 60×1.40-NA lens). Images captured 3 or 15 min after anaphase onset were analysed. To generate the average intensity profiles of fluorescently tagged tubulin or MAP65c–citrine perpendicular to the cell plate, the captured images were rotated for the vertical orientation of the division plane. The centre of the intensity distribution in each image was manually aligned based on the tubulin intensity peak or the MAP65c signal. Specific square areas (Fig. 3f and 3g; 21 × 5.2 µm, Fig. 3i; 16.5 × 16.5 µm, Extended Fig. 5d; 16.5 × 16.5 µm, Extended Fig. 5e; 16.5 × 8.2 µm) were cropped from the rotated image, and mean intensity of each vertical pixel column was calculated and plotted. The profiles were normalised to the brightest pixels.

#### Mean square displacement

Spinning disc confocal fluorescence time-lapse imaging (100×1.45-NA lens) of protonemal cells expressing SCAMP4–mNG was performed every 1.2 s, and single-plane images were analysed. SCAMP4 signals in the phragmoplast were manually tracked using FIJI software. The calculation of the MSD, statistical analysis, fitting, and plotting of the data were performed using self-built Python codes. MSD as a function of time-lag *Δt* was calculated as below:

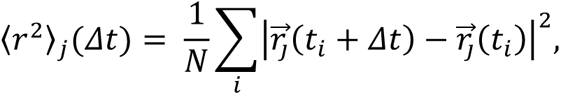

where the suffixes *i* and *j* are time point and trajectory, respectively, *Δt* is given time-lag, 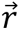 is the position vector of a particle, *t*_*i*_ is *i*-th time points of a trajectory, and *N* is the total number of *i* included in the calculation (not the total number of time points in a trajectory). The medians and means of the MSD were calculated over the trajectories. For statistical comparisons of medians of MSD between the two groups, p-values at each *Δt* were estimated using 500,000 resampled data reconstructed by bootstrapping and adjusted by the Holm method (α = 0.05). MSD was fitted using the following formula:

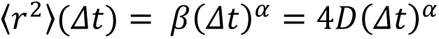

using the scipy.optimize.curve.fit function. SD values in the estimated parameters were computed using the square root of the diagonal components of the approximate covariance of the parameters.

#### Velocity and run length in vitro

TIRF time-lapse images (100×1.49-NA lens) were taken every 1–3 s. In the gliding assay, microtubules were manually selected and analysed by generating kymographs. In the single-motility assay, microtubules on which single or multiple fluorescent Kinesin12-II particles were continuously moving were manually selected and analysed by generating kymographs. The velocity and run length were obtained from the slopes of particle movements in the kymographs.

#### Statistics

Statistical analyses were performed using R version 4.3.1. The Shapiro–Wilk test was used to ascertain the normality of the data distribution across all samples. To evaluate the homogeneity of variances, an *F*-test was conducted for comparisons involving two groups, whereas Bartlett’s test was used for analyses involving multiple groups. If the samples (comprising two groups) had both a normal distribution and equal variances, Student’s *t*-test was used. If the samples (comprising two groups) had a normal distribution, but not equal variance, Welch’s two-sample *t*-test was used. If the samples (comprising two groups) did not have a normal distribution but had equal variance, Mann–Whitney *U* test was applied. If the samples (comprising two groups) had neither a normal distribution nor equal variance, Brunner– Munzel test was used. If the samples (multiple groups) did not have a normal distribution, they were analysed using the Aligned Rank Transform (ART) ANOVA with ARTool. Significant results were further analysed applying Tukey’s test. The statistical significance of the data was denoted using the following conventions for p-values: * < 0.05, ** < 0.01, *** < 0.001 and **** < 0.0001. Data from multiple experiments were pooled because of insufficient sample sizes in individual experiments. All experiments were performed independently at least twice.

### Accession numbers

*Physcomitrium patens Kinesin12-IIa*; Pp3c7_19581, *Kinesin12-IIb*; Pp3c7_19550, *Kinesin12-IIc*; Pp3c9_810, *Kinesin12-IId*; Pp3c17_5100, *SCAMP4*; Pp3c1_27370, *CalS5*; Pp3c9_4140, *Map65c;* Pp3c2_22900, *EB1b*; Pp3c20_6890.

## Acknowledgements

We are grateful to Gohta Goshima, Elena Kozgunova, Takema Sasaki and Yoshihisa Oda for critical reading of the manuscript; Peishan Yi for the plasmids; and Maya Hakozaki, Kyoko Kawamura, Chiemi Koketsu, Noiri Oguri and Rie Inaba for technical assistance. This work was funded by the JSPS KAKENHI (22K15138 to M.Y.) and a Naito Grant for female scientists after maternity leave (to M.Y.).

## Author contributions

Conceptualisation, M. Y.; investigation, M. Y.; formal analysis, M. Y. and H. J. M.; methodology, M. Y. and H. J. M.; writing, M. Y.; funding acquisition, M. Y.

## Movie legends

### Supplementary Video 1

SCAMP4–mNG dynamics in the phragmoplast and the cytoplasm of dividing protonemal tip cells. The yellow and red arrowheads indicate directional and diffusive movements of SCMAP4–mNG, respectively. Videos were acquired using spinning-disc confocal microscopy and a single plane is presented. Faint chloroplast autofluorescence was visualised.

### Supplementary Video 2

Cell division in the control and the *kin12-IIabcd-2* mutant. Dividing protonemal tip cells expressing GFP–α-tubulin (green) and histoneH2B–mCherry (magenta) were imaged with spinning-disc confocal microscopy, and maximum intensity projected images (2 µm × 3 sections) are presented.

### Supplementary Video 3

Map65 localisation during cytokinesis. Dividing protonemal tip cells expressing mCherry–α-tubulin (magenta) and Map65c–citrine (green). The control and the *kin12-IIabcd-1* mutant were imaged using spinning-disc confocal microscopy, and a single plane is presented.

### Supplementary Video 4

Dynamics of EB1 proteins in phragmoplasts. Dividing protonemal tip cells expressing EB1b–mNG (green) in the control and the *kin12-IIabcd-1* mutant were imaged using spinning-disc confocal microscopy, and a single plane is presented.

### Supplementary Video 5

Division of BFA-treated cells. Dividing protonemal tip cells expressing mCherry–α-tubulin (magenta) and SCAMP4–citrine (green), treated with either 0.5% DMSO or 50 µM BFA, were imaged with spinning-disc confocal microscopy, and a single plane is presented. The chemicals were applied to the samples 15–30 min before imaging.

### Supplementary Video 6

Accumulation of cell plate materials. Dividing protonemal tip cells expressing mCherry–α-tubulin (magenta) and SCAMP4–citrine (green) in the control, *kin12-IIabcd-1* mutant, and *kin12-IIabcd-1*/Kin12-IIc were imaged with spinning-disc confocal microscopy, and a single plane is presented.

### Supplementary Video 7

CalS5 dynamics during cell division. Dividing protonemal tip cells expressing mCherry–α-tubulin (magenta) and mNG–CalS5(green) in the control and the *kin12-IIabcd-1* mutant were imaged with spinning-disc confocal microscopy, and a single plane is presented.

### Supplementary Video 8

Dynamics of SCAMP4 in phragmoplasts. Dividing protonemal tip cells expressing SCMAP4 in the control and the *kin12-IIabcd-1* mutant were imaged using spinning-disc confocal microscopy, and a single plane is presented. Arrowheads indicate the directional movements of SCAMP4–mNG.

### Supplementary Video 9

In vitro single-molecule motility assay using purified Kin12-IIc^motor^–sfGFP. Microtubules (magenta) attached to glass and Kin12-IIc^motor^–sfGFP (green) were imaged using total internal reflection microscopy. The arrowhead and arrow indicate directional and diffusive movements, respectively.

### Supplementary Video 10

Movement of Kin12-IIc–mNG in the endoplasm. The endoplasmic microtubules of a protonemal cell expressing mCherry–α-tubulin (magenta) and Kin12-IIc–mNG (green) was imaged using oblique illumination fluorescence microscopy. Chloroplast autofluorescence is also faintly visualised in the green channel.

### Supplementary Video 11

Localisation of Kin12-IIc truncated proteins during cytokinesis. Dividing protonemal tip cells expressing mCherry–α-tubulin (magenta) and Kin12-IIc truncation variants tagged with mNG (green) in the *kin12-IIabcd-1* mutant were imaged with spinning-disc confocal microscopy, and a single plane is presented. Chloroplast autofluorescence is also faintly visualised in the green channel.

### Supplementary Video 12

Dynamics of the C-terminus truncation during cytokinesis. Dividing protonemal tip cells expressing mCherry–α-tubulin (magenta) and Kin12-IIc truncation (1022–1345 aa) tagged with mNG (green) in the control and the *kin12-IIabcd-1* mutant were imaged with spinning-disc confocal microscopy, and a single plane is presented. Chloroplast autofluorescence is also faintly visualised in the green channel.

### Supplementary Video 13

Development of the early gametophores. Dividing gametophore initial cells expressing GFP–α-tubulin (green) and histoneH2B–mCherry (magenta) in the control, *kin12-IIabcd-2* mutant and *kin12-IIabcd-2*/Kin12-IIc–cerulean were imaged using spinning-disc confocal microscopy. Maximum intensity projected images (2 µm × 21 sections) are presented.

## Tables

**Supplementary Table 1.**
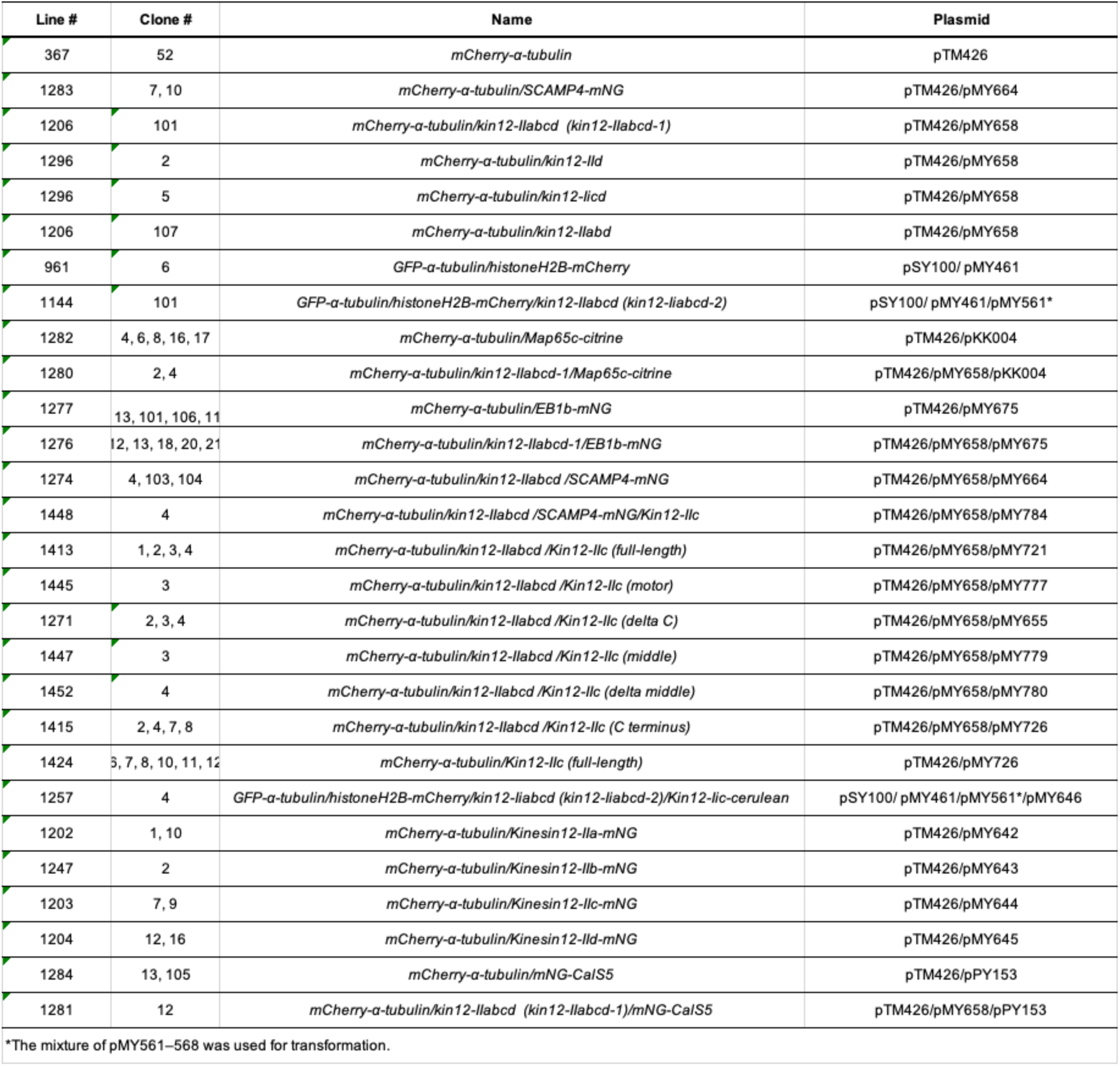
Moss strains used in this study.

**Supplementary Table 2.** Plasmids used in this study.

| Plasmid # | Description | Accession number | Targeting locus | *P. patens* promoter | *P. patens* resistance |
| --- | --- | --- | --- | --- | --- |
| pMY461 | PpH2B-mCherry integrant | Pp3c13_23540 | endogenous | - | - |
| pMY561 | Kinesin12-IIa sg1 NTC | Pp3c7_19581 | - | U6 promoter | NTC |
| pMY562 | Kinesin12-IIa sg2 NTC | Pp3c7_19581 | - | U6 promoter | NTC |
| pMY563 | Kinesin12-IIb sg1 NTC | Pp3c7_19550 | - | U6 promoter | NTC |
| pMY564 | Kinesin12-IIb sg2 NTC | Pp3c7_19550 | - | U6 promoter | NTC |
| pMY565 | Kinesin12-IIc sg1 NTC | Pp3c9_810 | - | U6 promoter | NTC |
| pMY566 | Kinesin12-IIc sg2 NTC | Pp3c9_810 | - | U6 promoter | NTC |
| pMY567 | Kinesin12-IId sg1 NTC | Pp3c17_5100 | - | U6 promoter | NTC |
| pMY568 | Kinesin12-IId sg2 NTC | Pp3c17_5100 | - | U6 promoter | NTC |
| pMY642 | Kin12IIa-mNG_G418 knockin | Pp3c7_19581 | endogenous | - | G418 |
| pMY643 | Kin12IIb-mNG_G418 knockin | Pp3c7_19550 | endogenous | - | G418 |
| pMY644 | Kin12IIc-mNG_G418 knockin | Pp3c9_810 | endogenous | - | G418 |
| pMY645 | Kin12IId-mNG_G418 knockin | Pp3c17_5100 | endogenous | - | G418 |
| pMY646 | EF1a::Kinesin12-IIc Cerulean PTA1_Hyg | Pp3c9_810 | PTA1 | EF1a | Hyg |
| pMY652 | EF1a::Ppkin12IIc rigor -mNG Hyg HB7 | Pp3c9_810 | HB7 | EF1a | Hyg |
| pMY655 | EF1a::Ppkin12IIc tail less 2-mNG Hyg HB7 | Pp3c9_810 | HB7 | EF1a | Hyg |
| pMY658 | Kin12II abcd multi sg NTC (pMY561, 565, 567) |  | - | U6 promoter | NTC |
| pMY664 | SCAMP4-mNG KI G418 | Pp3c1_27370 | endogenous | - | G418 |
| pMY675 | EB1b-mNG integrant G418 | Pp3c20_6890 | endogenous | - | G418 |
| pMY721 | kinesin12IIc::kinesin12IIc(1-1346)-mNG Hyg | Pp3c9_810 | HB7 | endogenous | Hyg |
| pMY726 | kinesin12IIc::kin12IIc(1022-1346)-mNG Hyg | Pp3c9_810 | HB7 | endogenous | Hyg |
| pMY742 | Kin12IIc(123-523)-sfGFP-6xHis bacterial expression | Pp3c9_810 | - | - | - |
| pMY777 | kinesin12IIc::Kin12IIc(1-523)-mNG Hyg | Pp3c9_810 | HB7 | endogenous | Hyg |
| pMY778 | kinesin12IIc::mNG-Kin12IIc-headless(473-1346)-FLAG | Pp3c9_810 | HB7 | endogenous | Hyg |
| pMY779 | kin12IIc::kin12IIc(473-850)-mNG | Pp3c9_810 | HB7 | endogenous | Hyg |
| pMY780 | kin12IIc::kin12IIc(1-523+1022-1346)-mNG | Pp3c9_810 | HB7 | endogenous | Hyg |
| pMY784 | Kinesin12IIc::Kin12IIc-FLAG_Hyg_HB7 | Pp3c9_810 | HB7 | endogenous | Hyg |
| pKK004 | Map65c-citrine knockin | Pp3c2_22900 | endogenous | - | G418 |
| pPY153 | mNG-CalS5 knockin | Pp3c9_4140 | endogenous | - | - |

**Supplementary Table 3.**
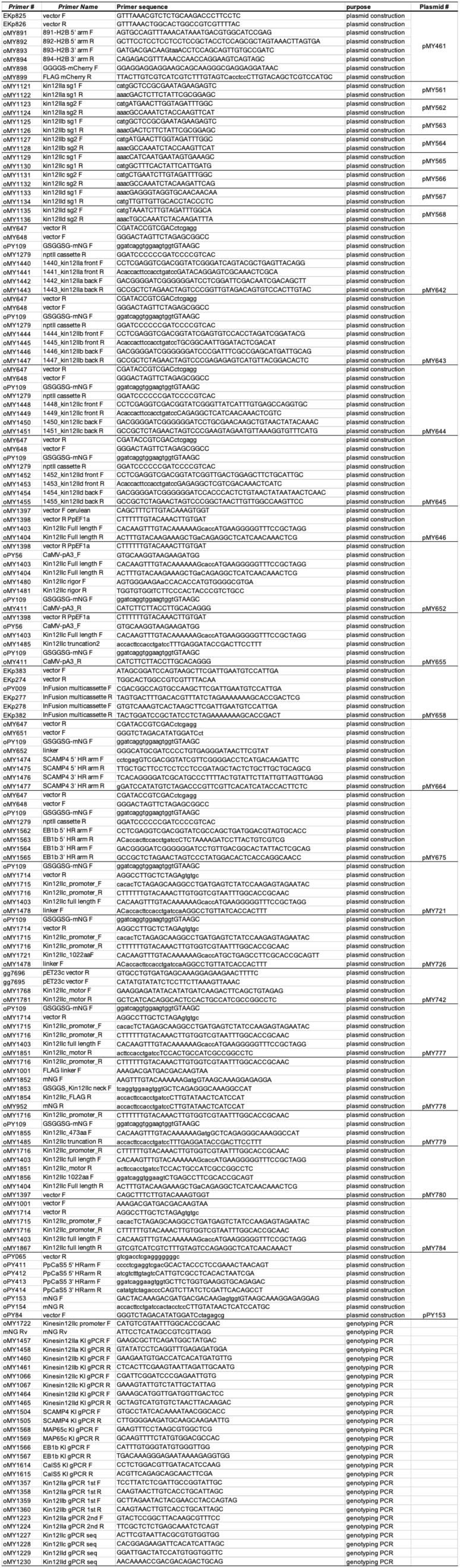
Primers used in this study.

**Supplementary Table 4.**
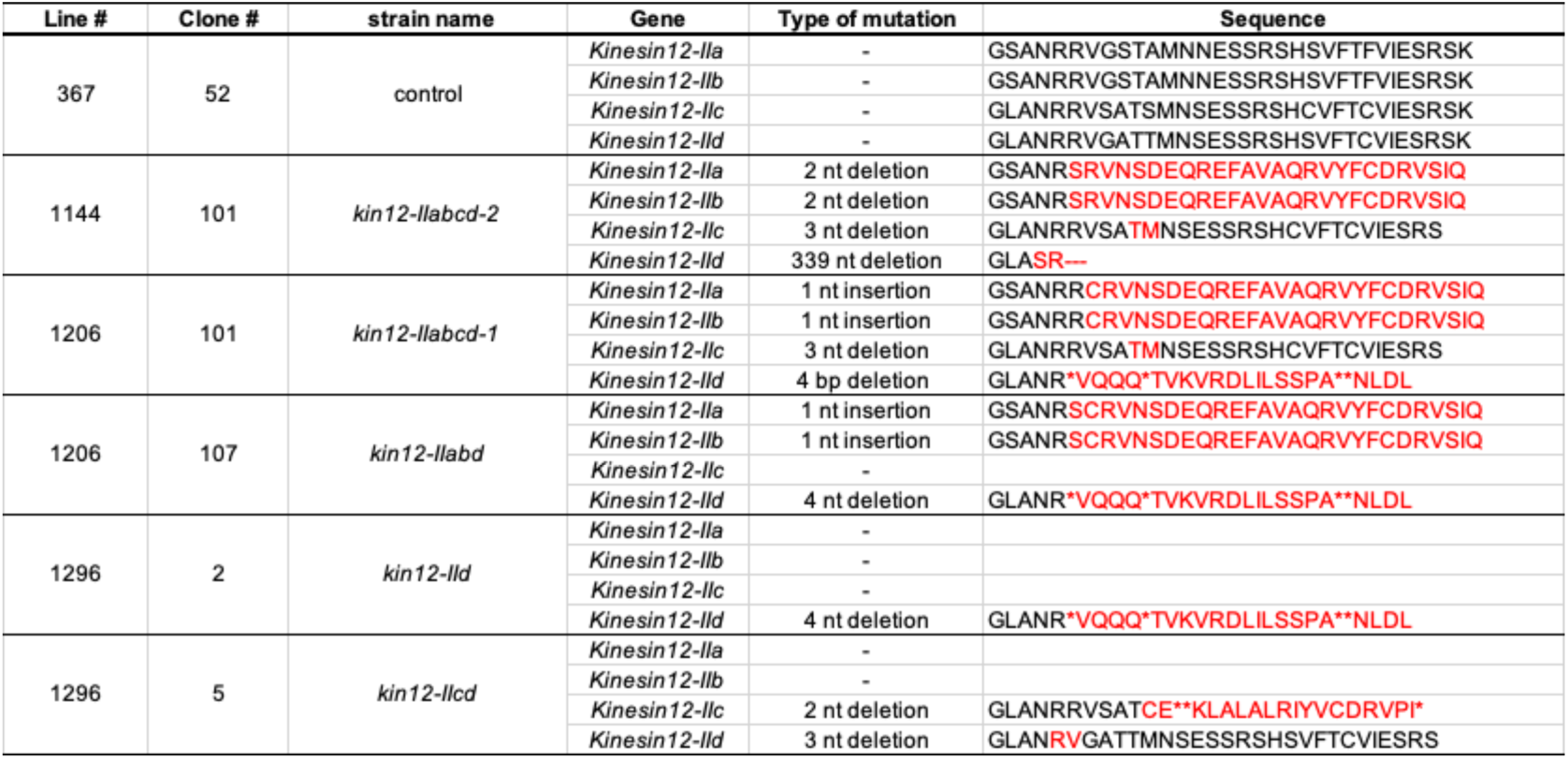
Amino acid sequences of the mutants used in this study.

**Supplementary Fig. 1:**
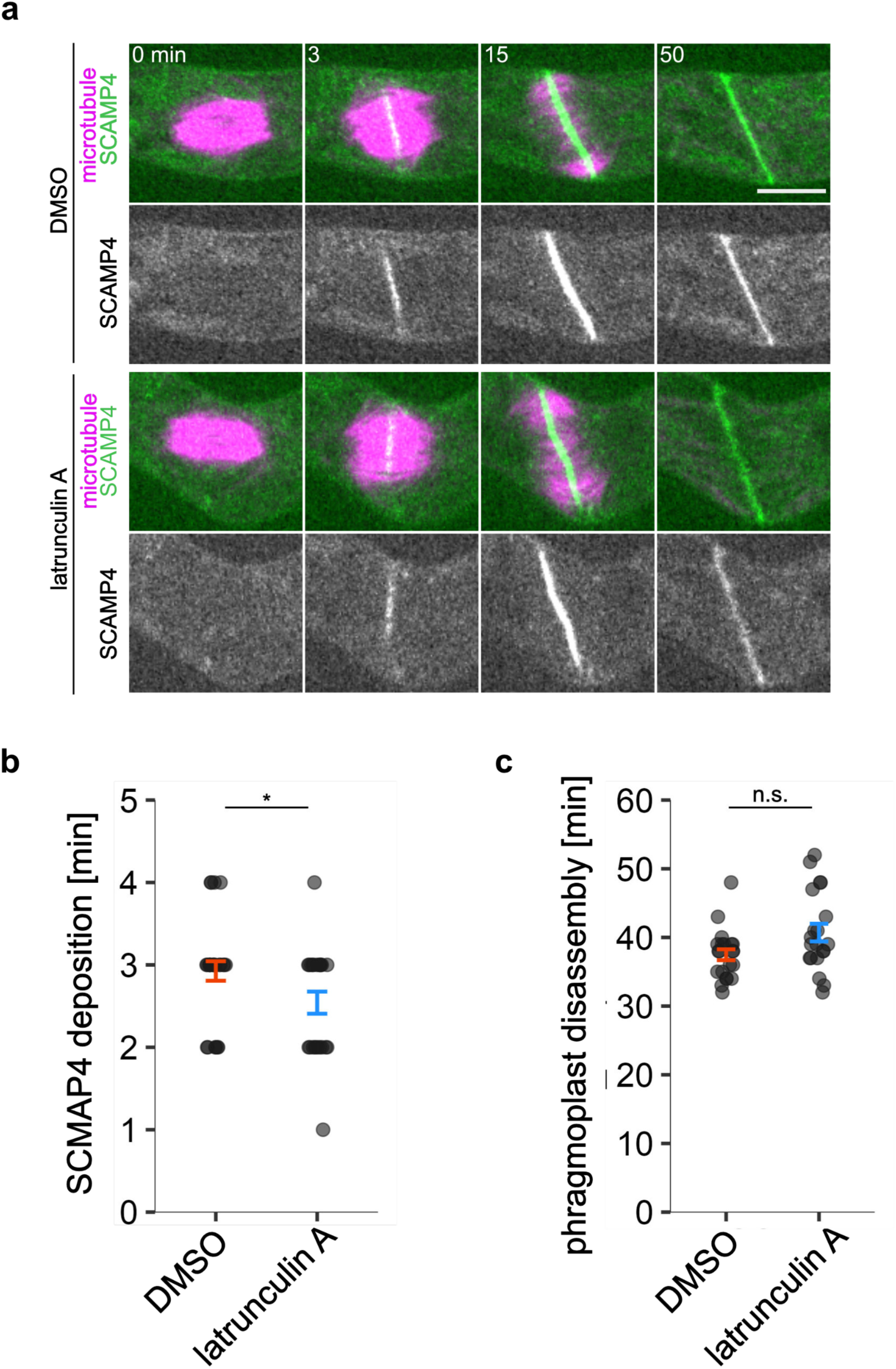
Actin is dispensable for cell plate material accumulation in moss protonemal cells. (a) Snapshots of a dividing protonemal tip cell expressing mCherry–α-tubulin (magenta) and SCAMP4– mNG (green). 0.5% DMSO or 5 µM latrunculin A were applied to the sample before imaging acquisition. A single plane of confocal images is presented. Time 0 corresponds to anaphase onset. Chloroplast autofluorescence is also faintly visualised in the green channel. Scale bar, 10 µm. (b, c) Comparison of the duration of cell plate deposition and phragmoplast disassembly. (b) DMSO, 2.9 ± 0.1 min (mean ± SEM), n = 27; latrunculin A, 2.5 ± 0.1 min, n = 24, *P* value was calculated using Mann-Whitney *U* test, *P* < 0.05. (c) DMSO, 37.5 ± 0.8 min (mean ± SEM), n = 21; latrunculin A, 40.7 ± 1.3 min, n = 20, *P* value was calculated using Mann–Whitney *U* test, *P* > 0.05.

**Supplementary Fig. 2:**
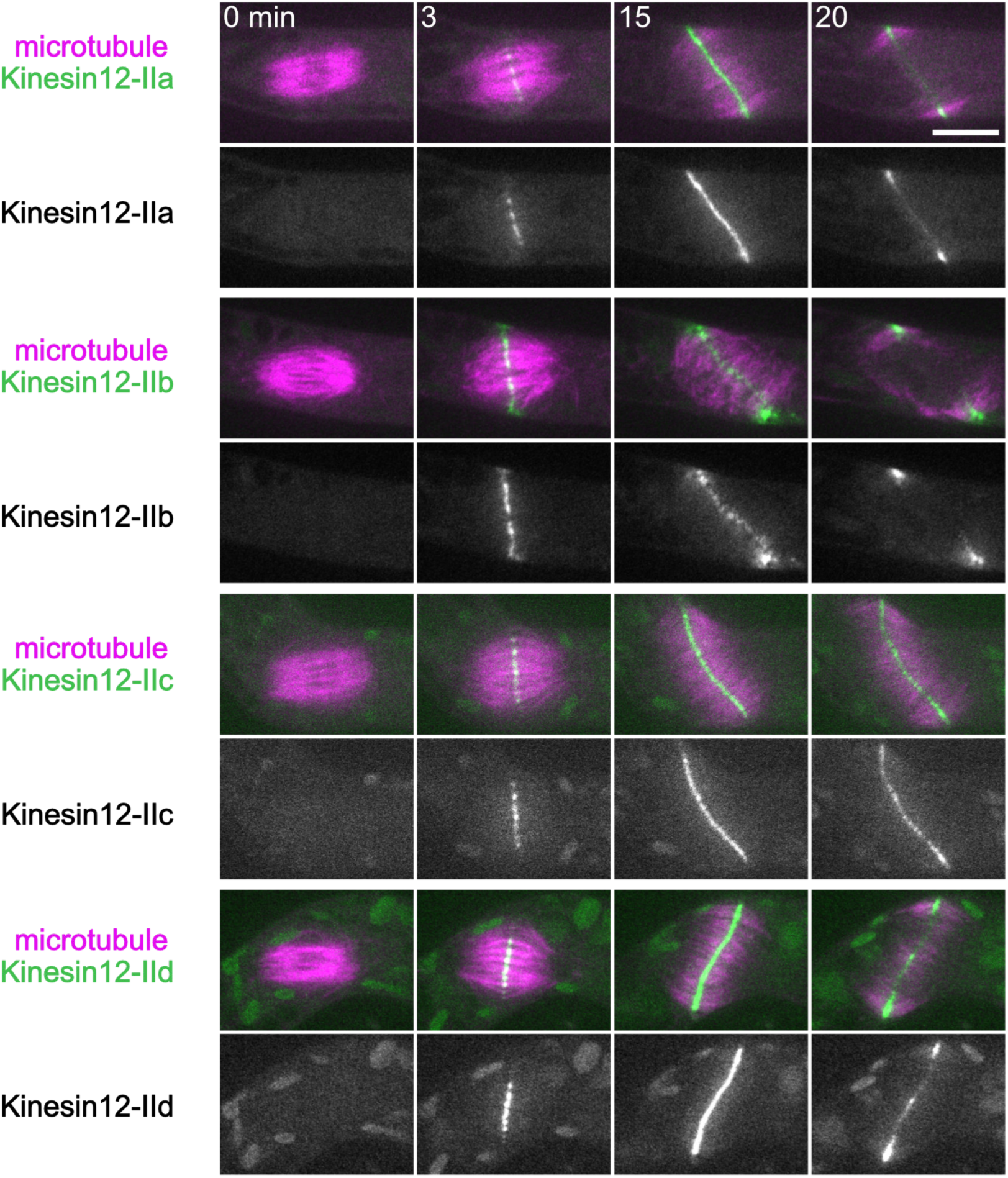
Kinesin12-II paralogues localise in the phragmoplast during cytokinesis. Snapshots of a dividing protonemal tip cell expressing mCherry–α-tubulin (magenta) and Kinesin12-II proteins tagged with mNG (green). A single plane of confocal images is presented. Time 0 corresponds to anaphase onset. Chloroplast autofluorescence is also faintly visualised in the green channel. Scale bar, 10 µm.

**Supplementary Fig. 3:**
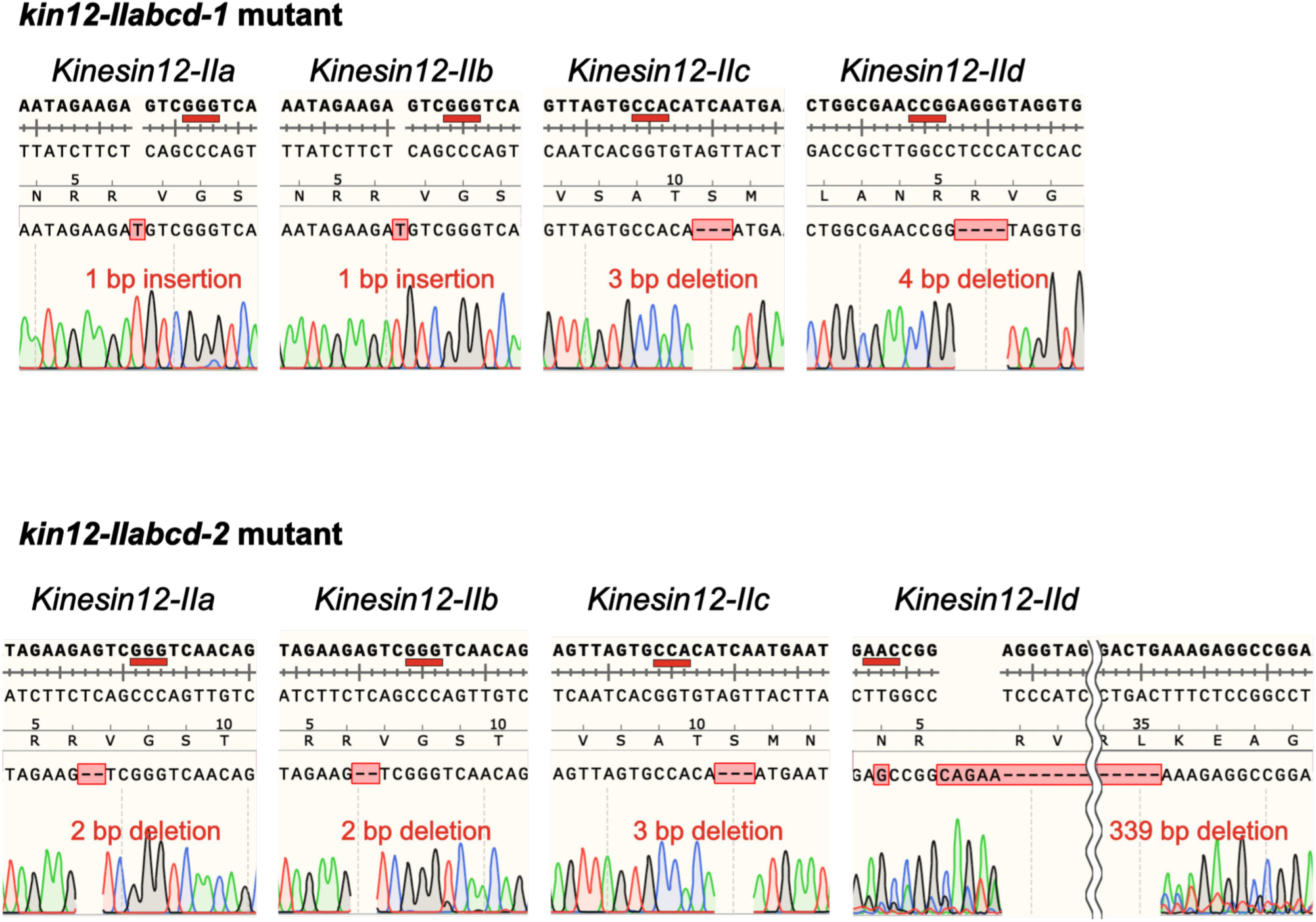
*Kinesin12-II* genes are disrupted by CRISPR mediated genome editing in the quadruple mutants. Sequencing data showing frameshift mutations and amino acid deletions in the *kin12-IIacbd-1* and *kin12-IIabcd-2* mutants. The SnapGene sequence files are presented with some modifications.

**Supplementary Fig. 4:**
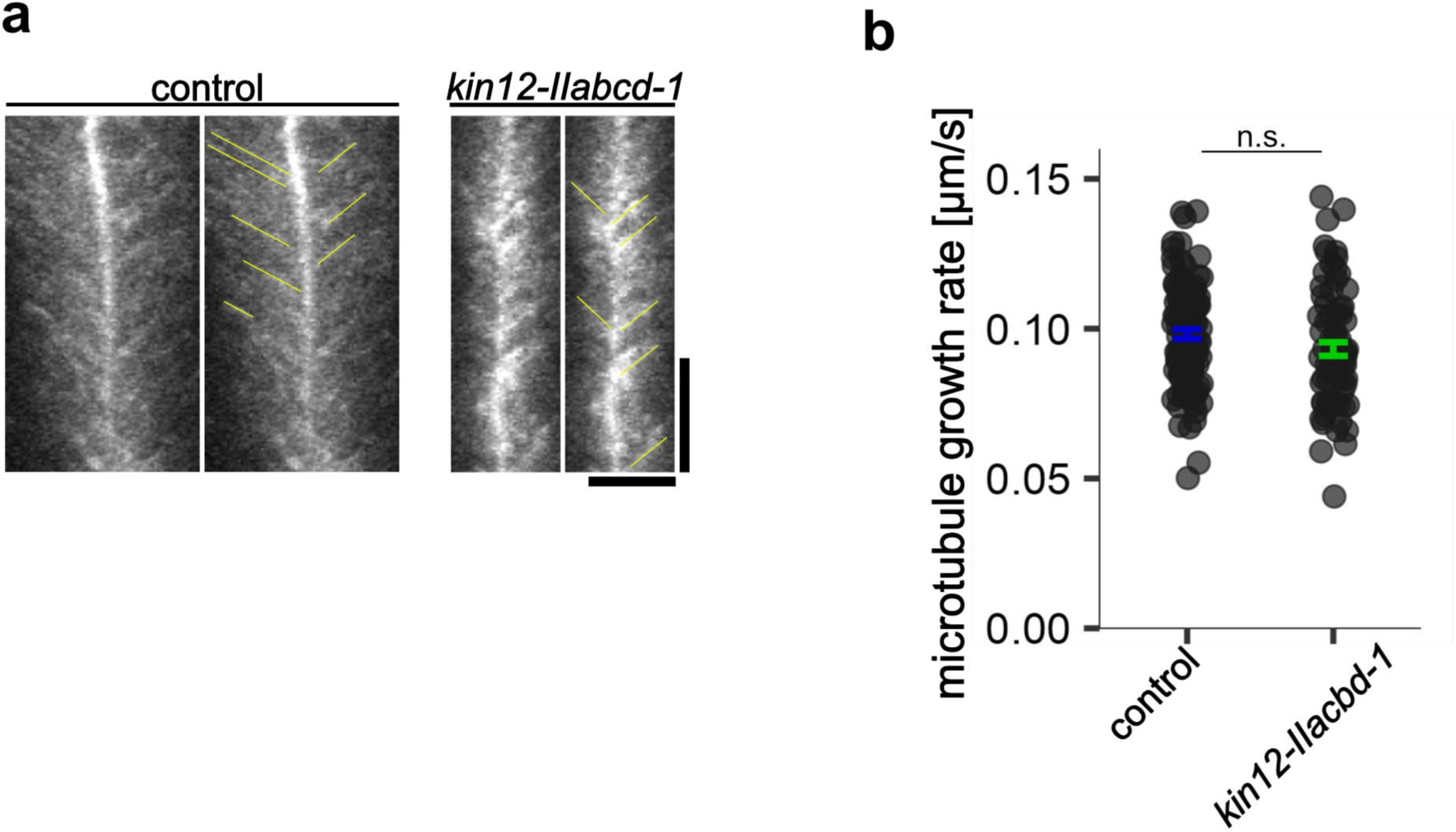
Microtubule polarity and growth rate are not altered by *Kinesin12-II*disruption. (a) Kymographs showing EB1b–mNG movement in the phragmoplast. Kymographs were generated along the phragmoplast microtubules, and the strong signal centred in the kymograph shows the midzone position. Horizontal bar, 5 µm; vertical bar, 100 s. (b) Comparison of the EB1-mNG velocity moving towards the midzone in the phragmoplast. control, 0.098 ± 0.002 µm/s (mean ± SEM), n = 114; *kin12-IIabcd-1*, 0.093 ± 0.002 µm/s, n = 77, *P* value was calculated using student *t*-test, *P* > 0.06.

**Supplementary Fig. 5:**
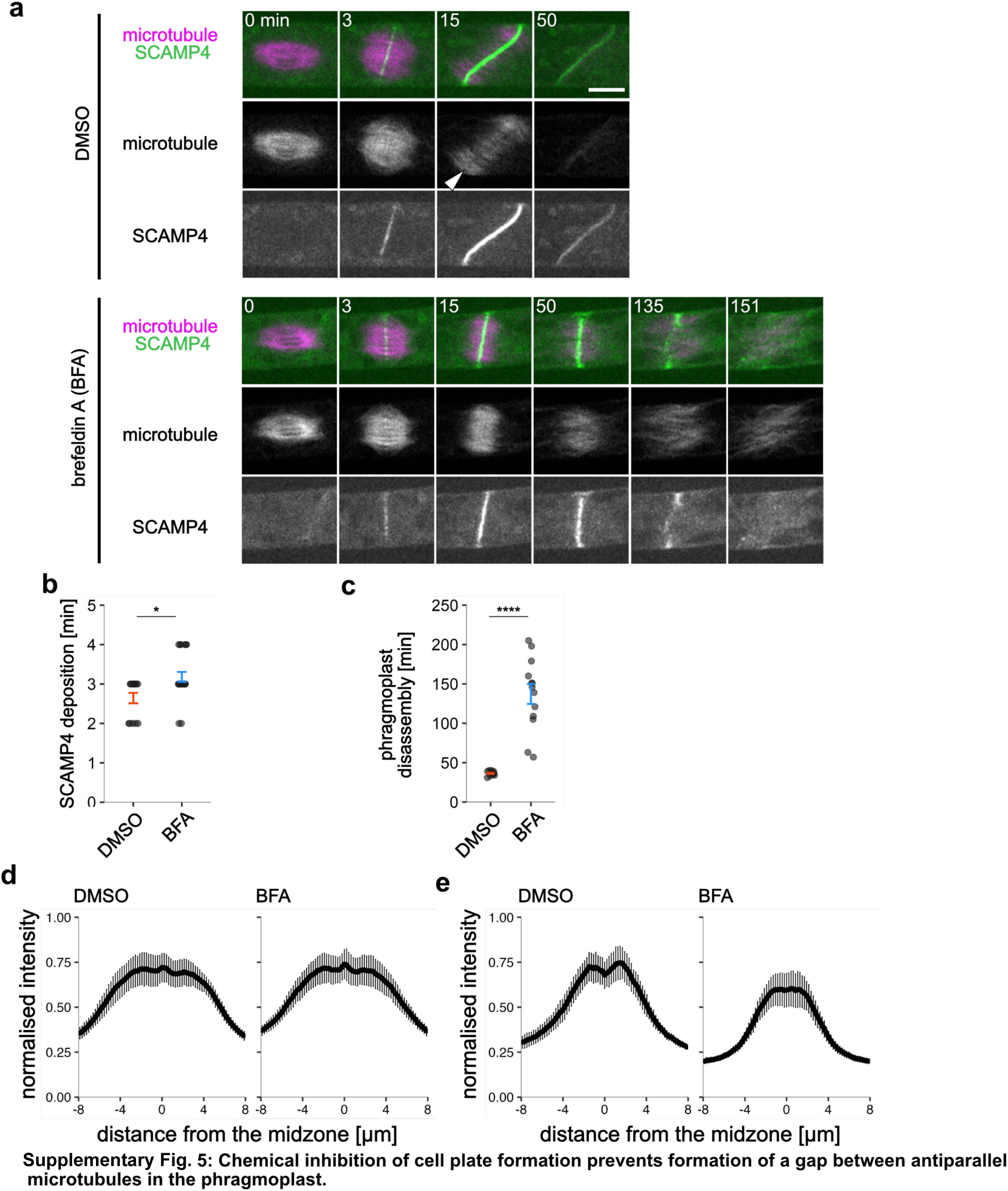
Chemical inhibition of cell plate formation prevents formation of a gap between antiparallel microtubules in the phragmoplast. (a) Snapshots of a dividing protonemal tip cell expressing mCherry–a-tubulin (magenta) and SCAMP4– mNG (green). 0.5% DMSO or 50 µM Brefeldin A was applied to the sample 15–30 min prior to imaging acquisition. A single plane of confocal images is presented. The white arrowhead indicates the position of the midzone gap in the control cell. Time 0 corresponds to anaphase onset. Chloroplast autofluorescence is also faintly visualised in the green channel. Scale bar, 10 µm. (b, c) Comparison of the duration of cell plate deposition and phragmoplast disassembly after BFA treatment. (b) DMSO, 2.6 ± 0.1 min (mean ± SEM), n = 14; BFA, 3.2 ± 0.1 min, n = 22. *P* value was calculated using Mann-Whitney *U* test, *P* < 0.05; (c) DMSO, 36.4 ± 0.8 min (mean ± SEM), n = 13; BFA, 137.2 ± 12.7 min, n = 13. *P* value was calculated using Welch’s *t*-test, *P* < 0.0001. (d, e) Normalised intensity profile of mCherry–a-tubulin in the phragmoplast of the control treatment (left) and the BFA treatment (right). The intensity was measured perpendicular to the division plane. The midzone position was plotted at 0 µm in the graph. The mean intensity is indicated by a black line, and the error bars indicate the SD values. The phragmoplast was analysed 3 min after anaphase onset in (d) and 15 min after anaphase onset in (e): (d), DMSO, n = 10 cells; BFA, n = 12 cells. (g), DMSO, n = 10 cells; BFA, n = 12 cells.

**Supplementary Fig. 6:**
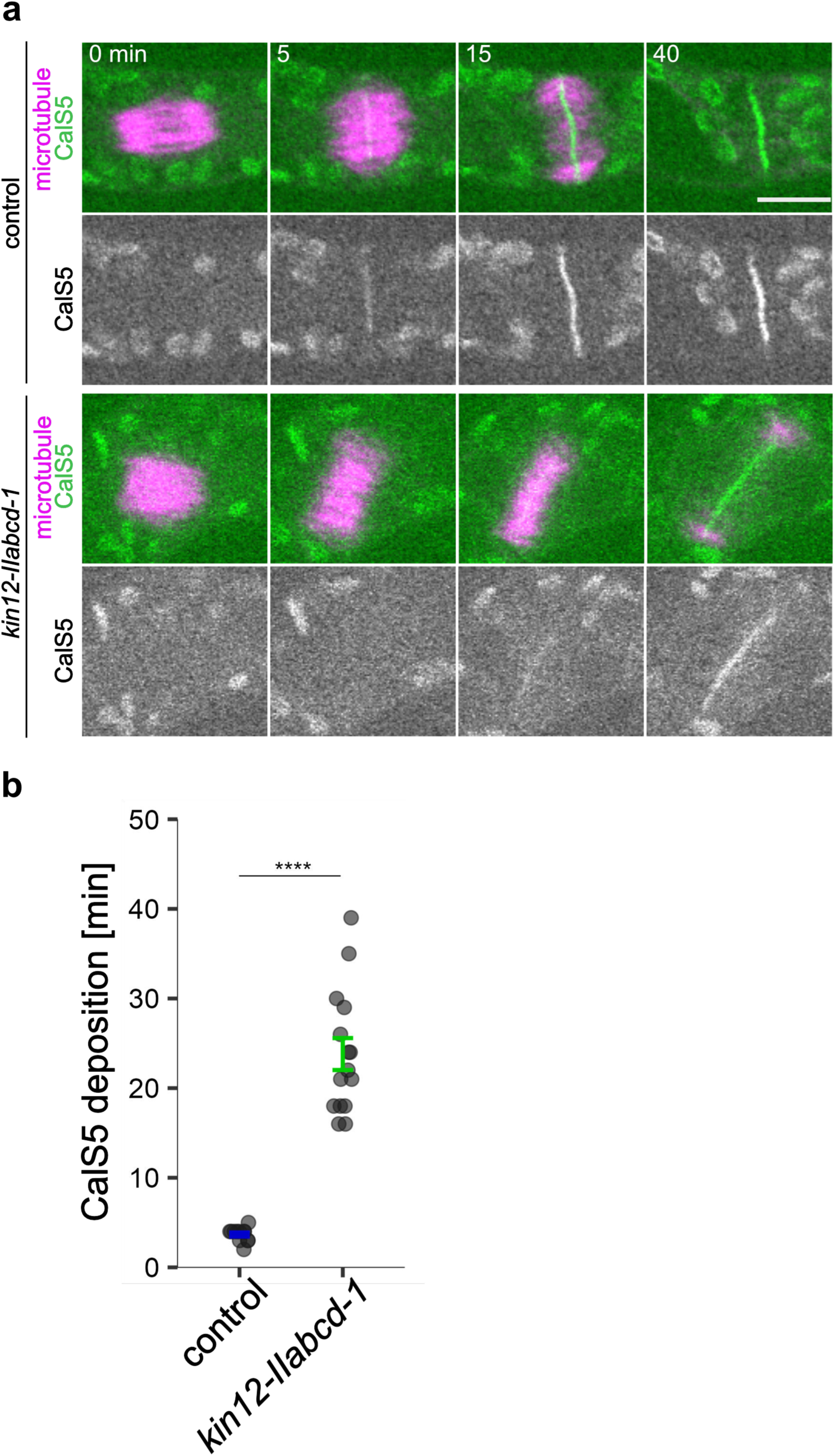
Deposition of CalS5 at the midzone is prolonged in the *kin12-IIabcd-1* mutant. (a) Snapshots of a dividing protonemal tip cell expressing mCherry–α-tubulin (magenta) and mNG– CalS5 (green) in the control and the *kin12-IIabcd-1* mutant. A single plane of confocal images is presented. Time 0 corresponds to anaphase onset. Chloroplast autofluorescence is also faintly visualised in the green channel. Scale bar, 10 µm. (b) Duration of cell plate deposition time detected by the mNG– CalS5 signal. Control, 3.7 ± 0.2 min (mean ± SEM), n = 12; *kin12-IIabcd-1*, 23.8 ± 1.8 min, n = 15, *P* value was calculated using Brunner–Munzel test, *P* < 0.0001.

**Supplementary Fig. 7:**
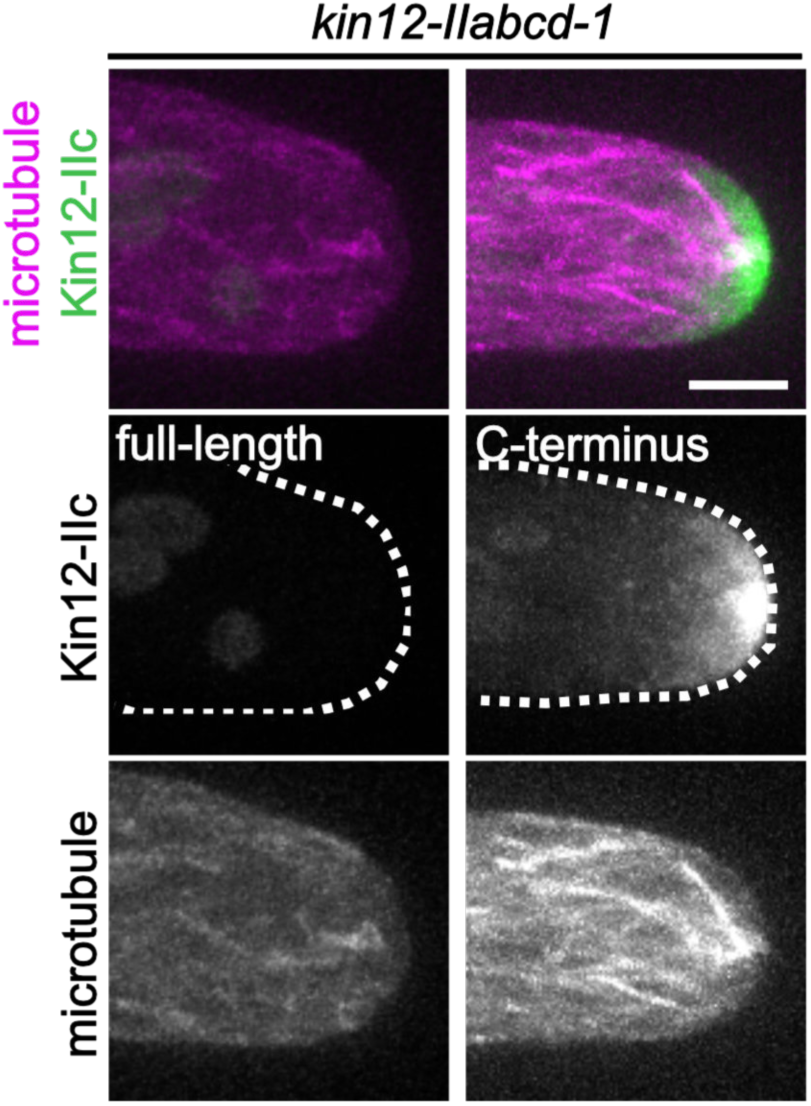
The C-terminus construct localised at the apex of the protonemal tip cell. Snapshots of a protonemal tip cell expressing mCherry–α-tubulin (magenta) and truncated Kin12-IIc-mNG (green). Maximum projected image of 9 confocal images acquired every 1µm is presented. Time 0 corresponds to anaphase onset. White dashed lines indicate cell edges. Chloroplast autofluorescence is also faintly visualised in the green channel. Scale bar, 5 µm.

